# MAP4 kinase-regulated reduced CLSTN1 expression in medulloblastoma is associated with increased invasiveness

**DOI:** 10.1101/2024.03.20.585872

**Authors:** Ece Sönmez, Shen Yan, Meng-Syuan Lin, Martin Baumgartner

## Abstract

De-regulated protein expression contributes to tumor growth and progression in medulloblastoma (MB), the most common malignant brain tumor in children. MB is associated with impaired differentiation of specific neural progenitors, suggesting that the deregulation of proteins involved in neural physiology could contribute to the transformed phenotype in MB.

Calsynthenin 1 (CLSTN1) is a neuronal protein involved in cell-cell interaction, vesicle trafficking, and synaptic signaling. We previously identified CLSTN1 as a putative target of the pro-invasive kinase MAP4K4, which we found to reduce CLSTN1 surface expression. Herein, we explored the expression and functional significance of CLSTN1 in MB. We found that *CLSTN1* expression is decreased in primary MB tumors compared to tumor-free cerebellum or brain tissues. CLSTN1 is expressed in laboratory-established MB cell lines, where it localized to the plasma membrane, intracellular vesicular structures, and regions of cell-cell contact. The reduction of CLSTN1 expression significantly increased growth factor-driven invasiveness. Pharmacological inhibition of MAP4 kinases caused increased CLSTN1 expression and CLSTN1 accumulation in cell-cell contacts. Co-culture of tumor cells with astrocytes increased CLSTN1 localization in cell-cell contacts, which was further enhanced by MAP4K inhibition.

Our study revealed a repressive function of CLSTN1 in growth-factor-driven invasiveness in MB and its potential implication in homologous and heterologous interactions of the tumor cells with cells in their microenvironment.

## Introduction

Medulloblastoma (MB) is the most common malignant pediatric brain tumor; it is subgrouped into WNT, SHH, Grp3, and Grp4 MB^1^. Despite considerably improved molecular diagnostics, prognosis remains dismal for patients with disseminated disease^2^. A commonality of MB across subgroups is impaired differentiation of specific neural progenitors as a mechanism associated with tumor initiation^3–8^. Neuronal-like, undifferentiated, and differentiating malignant populations of cells are the originating cells of WNT, SHH, and Group 3 tumors^4^. SHH tumors closely resembled granule neurons of varying differentiation states that correlated with patient age, whereas Grp3 tumors exhibit a developmental trajectory from primitive progenitor-like to more mature neuronal-like cells^4^. It follows that common to all MB are likely some neuronal traits, which may include neuron-specific Ca^2+^ signal transmission as well as the capability to form synapse-like cell-cell interactions. Additionally, OLIG2-expressing glial cells are highly enriched in therapy-resistant and recurrent medulloblastomas and display highly up-regulated gene sets of the neuronal and astrocyte lineages^5^. However, the contribution of neural-specific proteins involved in cell-cell interaction regulation and Ca^2+^ signaling to MB growth and tissue invasion remains poorly understood.

In a previous surface proteomic analysis in medulloblastoma cells, we found that knock-down of the Ser/Thr kinase MAP4K4 increased the plasma membrane association of the kinesin 1 adaptor protein Calsyntenin-1 (CLSTN1)^9^. CLSTN1 was originally identified as a proteolytically cleaved protein in postsynaptic membranes of both excitatory and inhibitory neurons, and through its C-terminal Ca^2+^ binding capabilities, it was suggested to control synaptic Ca^2+^ signaling^10^. CLSTN2 and CLSTN3 were subsequently identified based on high sequence similarities, whereby CLSTN1 was found to be ubiquitously expressed in all CNS neurons and CLSTN2 and 3 showed highest expression in GABAergic neurons^11^. CLSTN1 binds to the light chain of kinesin 1 (KLC1) via two conserved L/M-E/D-W-D-D-S motifs and is involved in anterograde tubulovesicular transport in neurons^12,13^. CLSTN1 recruits KLC1 to the trans-Golgi network (TGN), and overexpression of CLSTN1 causes ER and Golgi stacking^14^. This phenotype is explained by a disturbed stoichiometry of the physiological interaction of two CLSTN1 C-termini with one KLC1 molecule, which mediates vesicle cargo coupling to kinesin motors for anterograde transport^13^. CLSTN1 binding to KLC1 can be negatively regulated by phosphorylation of S460 in KLC1^15^. The repression of ERK was shown to reduce S460 phosphorylation and increase CLSTN1-KLC1 binding, suggesting that CLSTN1 trafficking could be controlled by reversible KLC1 S460 phosphorylation downstream of MAPK signaling^15^. Increased KLC1 S460 phosphorylation was observed in Alzheimer’s disease (AD), which suggested that reduced CLSTN1-KCL1 interaction may contribute to AD onset or progression^16^. CLSTN1 controls N-methyl-D-aspartate (NMDA) receptor subunit trafficking in pyramidal neurons during postnatal development in mice, and CLSTN1 knock-out (KO) impaired neuronal arborization and maturation^17^. CLSTN1 also regulates the transport of Rab5-positive endosomes towards specific compartments in developing axons, and it was suggested that this directed transport contributes to axon branching^18^. CLSTN1 also restricts the surface expression of ICAM5 in cultured neurons, which is necessary for proper spine maturation in dendrites. Reduced CLSTN1 and increased ICAM5 membrane localization further correlate with excessive formation of filopodia-like spines and impaired spine formation^19^.

Collectively, these studies on CLSTN1 indicate a critical function of CLSTN1 in the anterograde transport of cargo from ER/Golgi toward specific peripheral compartments in neuronal cells. Little is known about CLSTN1 function in cancer. ESRP1-mediated exon 11 skip splicing of CLSTN1 has been implicated as an anti-metastatic mechanism in gastric cancer, whereby the alternatively spliced, shortened CLSTN1 restricts migration and invasion by stabilizing E-cadherin-β-catenin binding^20^. The RNA binding protein AKAP8 was also implicated in CLSTN1 exon 11 skip splicing, yielding a short CLSTN1 isoform involved in maintaining an epithelial state and restricting metastasis in breast cancer^21^.

MAP4K4 is overexpressed in MB tissue compared to normal cerebellum and contributes to tissue invasion downstream of growth factor signaling in MB^22,23^. However, how MAP4K4 is connected to CLSTN1 and whether an interaction of MAP4K4 and CLSTN1 could be functionally relevant in MB is not known. Here, we explored whether CLSTN1 could contribute to the cancerous phenotype in MB and whether MAP4 kinase modulation of CLSTN1 expression could constitute an alternative mechanism to exon 11 skip splicing control of migration and invasion.

## Materials and methods

### Cells and tissue culture

#### Cell lines used

All cell lines used are described in the supplementary methods file. All media, reagents, and solutions used were filtered using 0.45 µm filters. The standard cell culture media were prepared using 10% heat-inactivated Fetal Bovine Serum (FBS), 1% Penicillin Streptomycin (P/S), and 1% L-glutamine. Serum-free media (SFM) were supplemented with 1% P/S and 1% L-glutamine only.

#### Generation of stable cell lines

Stable MB cell lines over-expressing mNeonGreen(mNG)-tagged CLSTN1 (CLSTN1-mNG) and V5-tagged CLSTN1 (CLSTN1-V5) were generated using lentiviral transduction and drug selection. Lentivirus was produced in HEK293-T cells transfected with 4.5 µg pPAX2, 3 µg pVSVG and 7.5 µg of either pLV-Bsd-CMV-hCLSTN1-3xGGGGS-mNeonGreen or pLV-Bsd-CMV-hCLSTN1-3xGGGGS-V5. Medium was replaced after 18h and virus-containing supernatant was collected 30 h after transfection. The supernatant was centrifuged (300xg, 5 min) and filtered using a 5 µm pore filter. 1 ml of filtered supernatant was added to the recipient cells in 1 ml complete medium with 10 µg/ml polybrene. Filtered supernatants were stored at −80°C until further use.

2 days post-transduction, the MB cells were expanded, and the following day, antibiotic selection was started using 5 µg/ml blasticidine dissolved in fresh complete medium. The selection was continued for two weeks and the medium containing blasticidine was changed every second day. Ectopic protein expression levels of CLSTN1-mNG and CLSTN1-V5 were confirmed using IB or flow cytometry (FC) analysis, and uniformity of expression by microscopy imaging. UW228 LA-EGFP_mCherry-Nuc9 cells were produced by lentiviral transduction of UW228 with pLenti-LA-EGFP and Nuc9mCherry. Two days after transduction, cells were transferred to selection medium containing 2 ug/ml puromycin for at least three passages. LA-EGFP expression was confirmed by microscopy imaging.

#### Gene expression analyses in human primary samples

mRNA expression analysis across tumor samples and healthy tissues was performed using the R2 Genomic Analysis & Visualization Platform. The following control datasets were used: Diseased brain: GSE9770; Brain regions: GSE11882; Cerebellum (ages of postmortem donors in years: 25 (m), 38 (m), 39 (f), 30 (m), 35 (m), 52 (m), 50 (f), 48 (f), 53 (f), 23 (f)): GSE3526. The following tumor datasets were used: Kool – 62: GSE10327; Delattre – 57; Pfister – 73: GSE49243; Gilbertson _ 76: GSE37418; Pfister – 223: PubMed 28726821; Hsieh – 31: GSE67851; denBoer – 51: GSE74195; Cavalli – 763: GSE85217. For genomic analysis across datasets, the Affymetrix Human Genome U133 Plus 2.0 Arrays were used.

#### Spheroid invasion assay (SIA)

SIA was performed and quantified according to^24^. In brief: 24h after siRNA transfection, 2’500 cells were seeded per well in 100 µl of complete medium in a 96-Well Clear Round Bottom Ultra-Low Mount Microplate (Corning™ Costar™ Cat#7007) and incubated overnight at 37° C, 5% CO_2_ conditions. After 24h, formation of compact and uniform spheroids was confirmed by light microscopy. 70 μl medium was then removed and replaced with 70 μl of a solution containing 2.7 mg/ml PureCol® bovine collagen (CellSystems, Cat#5005). Polymerized collagen hydrogels were overlaid with 100 μl of SFM containing growth factors and/or inhibitors 2X concentrated. The cells were allowed to invade the collagen matrix for 24 h, after which they were stained with Hoechst (Sigma Aldrich, Cat#B2883, 1:2000) for 3–4 h. The image acquisition was performed using an Operetta CLS High-Content Analysis System (PerkinElmer, Cat# HH16000000) using the 405 nm channel. Spheroids and the invading cells were defined based on the fluorescence threshold using Harmony software. For each tumor cell nucleus, the distance from the center of the spheroid was calculated. The total distance of invasion from the center of the spheroid was calculated by summing up the individual invasion distances of each cell using Harmony (Perkin Elmer). The data was displayed and statistically analyzed using Prism 9.3.1 software (GraphPad).

#### Immunoblot (IB)

Cells were lysed using RIPA buffer supplemented with cOmplete™, Mini Protease Inhibitor Cocktail (Roche, Cat#11836153001) and phosphatase inhibitors PhosSTOP™ (Roche, Cat#4906837001), and cleared by centrifugation. Protein concentration was determined using the Pierce™ BCA Protein Assay Kit (Thermo Fisher Scientific, Cat# 23225). Gel electrophoresis was performed on 4–20% Mini-PROTEAN® TGX™ Precast Protein Gels (Bio-Rad Laboratories AG, Cat#4561094/4561096). Proteins were transferred to Trans-Blot Turbo 0.2 µm Nitrocellulose Transfer Membrane (Bio-Rad, Cat#1704158/1704159) and membranes blocked with 5% non-fat milk for 1 h. For primary antibodies used see supplementary file 1, table 2. HRP-linked secondary antibodies were used with Western Blotting Substrate (Thermo Fisher Scientific, Cat#32209) or SuperSignal™ West Femto Maximum Sensitivity Substrate (Thermo Fisher Scientific, Cat#34095). The integrated density of immuno-reactive bands was quantified using Image Lab (Version 5.2.1, Bio-Rad Laboratories).

#### Subcellular protein fractionation

6 × 10^5^ cells/well were seeded in 6 cm dishes and incubated at 37^°^ C. After 24 h, 3 ml of media were removed, and the cells were collected by scraping into the remaining medium. The cells were washed twice in PBS and the cellular contents from the dry cell pellets was fractionated using the subcellular protein fractionation kit for cultured cells (ThermoFisher Scientific, Cat# 78840) exactly according to the manufacturer’s protocol. This protocol enabled the stepwise extraction of the proteins from the cytoplasm (CEB), membranes (MEB), nuclear soluble (NEB), chromatin-bound regions (ChrB), and cytoskeleton (Ck). The subcellular protein extracts were analyzed by IB as described above.

#### siRNA transfection

1.5 × 10^5^ DAOY, UW228, and ONS-76 cells, or 3 × 10^5^ D425, HD-MBO3, and D283 cells were seeded per well in 6-well plates. After 24 h, cells were transfected with 6 nM siRNA mixed with OptiMEM reduced serum medium (Thermo Fisher Scientific, Cat# 31985062) using Lipofectamine RNAiMAX transfection reagent (Invitrogen, Cat# 13778075). 6h after adding the transfection mixture, the transfection media was replaced with complete fresh medium, and the cells were cultured for another 24 – 72 h for further analysis.

#### Flow cytometry (FC) analysis

The cells were collected on ice by gentle scraping. PBS-washed cells were fixed in 4% PFA in PBS for 20 min at room temperature (RT). Cells were then either processed immediately or stored overnight at 4°C before staining. Biotin-tagged anti-CLSTN1 antibody diluted (1:100) in FC buffer was used for CLSTN1 detection. PE-labelled IgG1 antibody (1:300) in PBS containing 2% FBS (FC buffer) was used as the negative control. Samples were incubated at 4°C in the dark for 1h, and then washed once with 300 µl FC buffer. Anti-CLSTN1 labeled samples were subsequently incubated with PE-coupled streptavidin diluted (1:200) in FC buffer at 4°C in the dark for 30 min and washed 3 × 5 min with 300 µl FC buffer at RT. All samples were then filtered into 5 ml polystyrene round-bottom tubes (Falcon/Brunschwig, Cat# 352235) and stored at 4°C in dark until data acquisition. Data was acquired using a BD Fortessa^TM^ Cell Analyzer, analyzed using FlowJo 10.8.0 software, and plotted using GraphPad Prism 9.3.1 software.

### Immunofluorescence analysis (IFA)

200 µl of a suspension of 4×10^3^ LA-EGFP mCherry-Nuc9 cells/ml were seeded per well in an ibidi 8-well plate and incubated overnight. Cells were then fixed in complete medium containing 4% PFA for 20 min at 37°C. Cells were washed 2 × 5 min in PBS, permeabilized with 0.1 % Triton X100 in PBS for 20 min at RT and washed again 3 × 5 min in PBS. Cells were blocked with PBS supplemented with 5% FBS for 1h at RT. Primary antibodies were diluted in PBS +5% FBS and incubated overnight at 4°C in a humidified chamber. Cells were then washed 3 × 5 min in PBS and nucleic acids stained with DAPI (1:5000) diluted in PBS for 5 min at RT. 1 drop of pre-warmed (37°C) DAKO Glycergel mounting medium (DAKO, Cat#C0563) was added to each well. Image acquisition was performed either on a NikonTi2 widefield or an SP8 Leica confocal microscope.

#### Tumor co-culture with immortalized normal human astrocyte (iNHA^25^)

2 × 10^3^ UW228 LA-EGFP mCherry-Nuc9 cells and 2 × 10^3^ iNHA or primary mouse cerebellar astrocytes were cultured together per well in an 8-well ibidi plate in complete DMEM medium supplemented with 1% sodium pyruvate. DMSO (control) or 500 nM prostetin/12K treatments were performed for 24h after which cells were fixed and stained fixed and stained with anti-CLSTN1 as described for regular and anti-GFAP as described for IFA.

### [Ca^+2^]_i_ detection using GcaMP6 calcium indicators

1.5 × 10^5^ GcAMP6-expressing UW228 cells were seeded per well in a 6-well plate and transfected with either si1-CLSTN1 or siRNA Ctrl 24h after seeding as described above. After 24 h, 1×10^4^ trasnfected cells were seeded per well in an 8-well ibidi plate in 200 µl complete DMEM. After 48 h, the medium was replaced with freshly prepared artificial cerebrospinal fluid (aCSF) solution without phenol red. Image acquisition was performed on a Nikon Ti2 widefield fluorescence microscope using a 20x objective. For each imaging series, 601 frames were recorded at 2-second intervals for 20 min. The image analysis was performed with Fiji 2.0.0 software (ImageJ2) and the frame rate was set up as 10 fps. Overlay images and calcium traces were generated with phyton programming.

### Live imaging of CLSTN1-mNG intracellular dynamics

6 × 10^3^ UW228 CLSTN1-mNG cells were seeded per well in an 8-well ibidi plate in 200 µl complete DMEM overnight. Cells were treated with 500 nM Prostetin or an equivalent volume of the solvent DMSO (control) and incubated for 24 h. 24 h after start of the treatment, the cells were monitored using Leica SP8 Confocal Microscope at 63X objective using the 488-laser line. Images were recorded every 30 seconds for 10 minutes. The analysis was performed using Fiji 2.0.0 software (ImageJ2).

#### Cell proliferation analysis

1.5 × 10^5^ UW228 cells were seeded in 6-well plate for siRNA transfection. After 24h, cells were transfected with siCtrl and si1-CLSTN. 24h after transfection, 5 × 10^4^ cells/well of each condition were seeded in 6-well plates and cultured overnight. Cells were counted 48h and 72h post-transfection using Trypan blue in combination with an automated cell counter (NanoEnTek).

#### Statistical analysis

Statistical analysis was performed using GraphPad Prism 9.3.1 software. Unpaired Student’s t-test was used to determine the statistical significance of the difference between two groups, and One-way ANOVA repeated measures test was used for multiple comparisons. Results with p-values ≤ 0.05 were considered as significant (* = P < 0.05, ** = P < 0.01, *** = P < 0.001, **** = P <0.0001, ns, not significant).

## Results

### Reduced expression of *CLSTN1* in medulloblastoma

The Calsyntenin (CLSTN) family of proteins consists of three members, CLSTN1, CLSTN2 and CLSTN3^11^, with relatively low protein sequence identity scores (CLSTN1:CLSTN2 = 51.81%, CLSTN1:CLSTN3 = 55.20%, CLSTN2:CLSTN3 = 49.02%). By surface proteome analysis, we only detected CLSTN1 in MB cells^9^ and found that CLSTN1 belonged to a small sent of proteins that we found to be upregulated by MAP4K4 depletion (Fig. 1A). To more broadly assess expression levels of the three *CLSTNs* across human MB samples in comparison to normal brain regions and the cerebellum, we analyzed *CLSTN1*, *CLSTN2*, and *CLSTN3* mRNA expression across five publicly available MB datasets as well as an atypical teratoid rhabdoid tumor (ATRT) and an ependymoma dataset (Fig. 1B). This analysis revealed significantly lower levels of *CLSTN1* and *CLSTN3* in the MB samples in comparison to normal brain regions or cerebellum samples. In contrast, *CLSTN2* expression is significantly increased in MB compared to the cerebellum controls. We next assessed differential expression of *CLSTNs* across the 12 MB subtypes using the Cavalli dataset^26^, which comprises 763 primary MB tumor samples (Fig. 1C). We observed some variations in the expression levels between the subtypes, with increased *CLSTN1* and *CLSTN3* in SHH-gamma and Grp3-gamma, and decreased *CLSTN1* and *CLSTN3* in SHH-delta and Grp3-delta samples. Interestingly, the expression levels of *CLSTN2* within each subgroup vary considerably more compared to *CLSTN1*, which may indicate a tighter control of *CLSTN1* transcription. The expression of *CLSTN1* and *CLSTN3* correlates negatively with *MAP4K4* expression, whereas *CLSTN2* correlates positively (Fig. 1D), further indicating differential regulation of *CLSTN1* and *CLSTN3* expression compared to *CLSTN2*. *CLSTN1* (Fig. 1E) and *CLSTN3* (Fig. S1B) expression increase sharply in the cerebellum after birth^27^, unlike *CLSTN2*, where expression drops at birth (Fig. S1A). Consistent with a potential coregulatory mechanism, expression of *MAP4K4* in the cerebellum also drops sharply after birth^28^. In the developing human cerebellum^29^, *CLSTN1* (Fig. 1F, S2) and *CLSTN3* (Fig. S1D, S2) are strongly expressed in granule cells. *CLSTN3* is additionally expressed in glutamatergic deep nuclei neurons and unipolar brush cells. In contrast, *CLSTN2* is highly expressed in Purkinje cells (Fig. S1C) and in a subset of granule cells. Interestingly, *MAP4K4* expression is lowest in granule cells, where *CLSTN1* and *CLSTN3* are highly expressed, and highest in oligodendrocytes, where *CLSTN1* and *CLSTN3* are not expressed (Fig. 1G).

**Figure 1:**
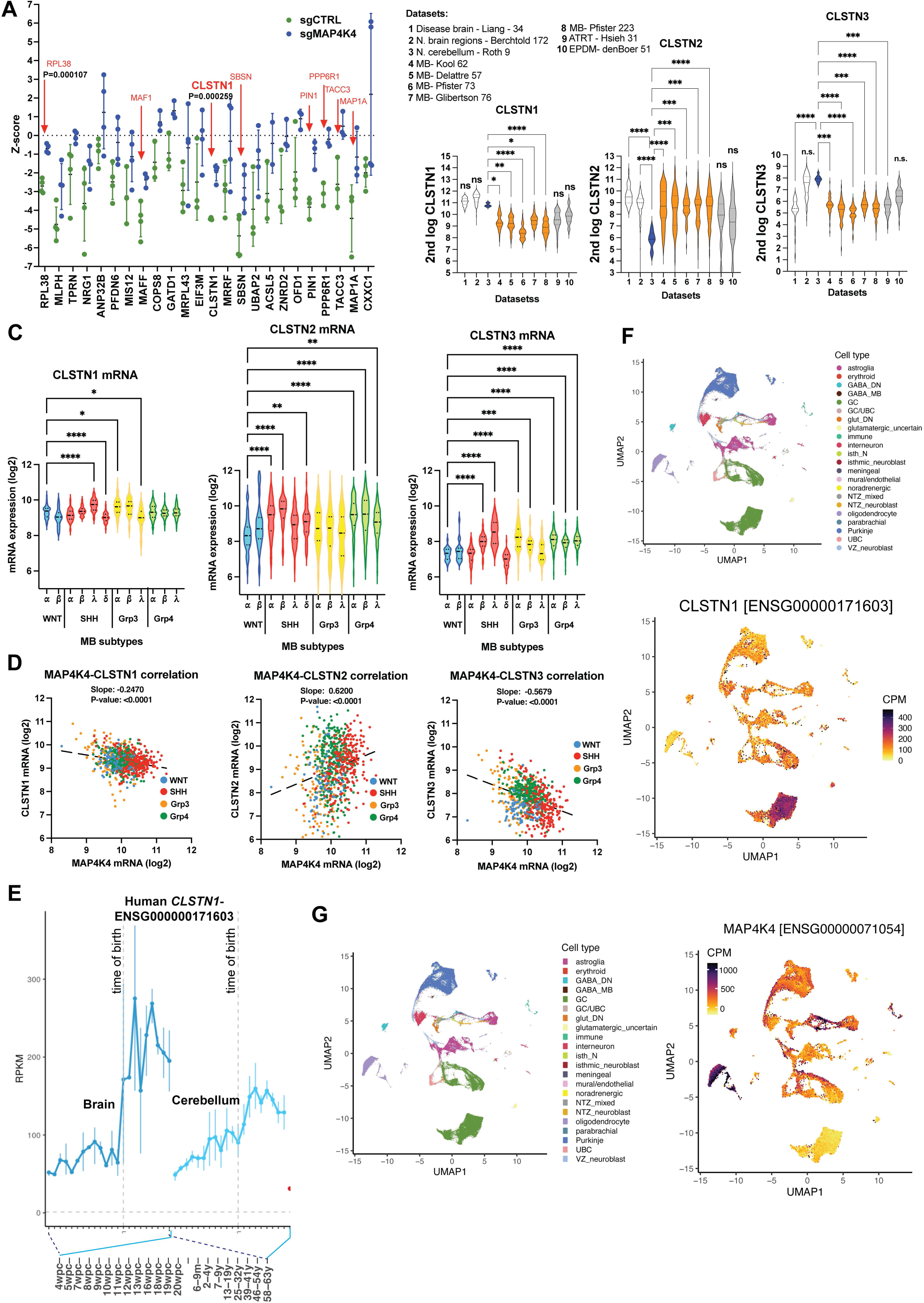
CLSTN1 expression is reduced in primary MB tumor samples. **A)** Differential protein expression in the plasma membrane-associated proteome between sgCTRL and sgMAP4K4 cells assessed by spatial mass-spectrometry analysis identified CLSTN1 as one of several specifically upregulated prooteins^30^. The mean Z-score and SD of up-regulated proteins in sgMAP4K4 cells are shown. n = 3 – 4 biological replicas. P values of multiple unpaired T-tests are shown when P=<0.05. **B)** mRNA expression analysis in three controls (diseased brain, normal brain regions and cerebellum), five primary MB (orange label), one ATRT, and one ependymoma dataset. Significant differences between CLSTN expression in tumor samples compared to cerebellum controls were calculated using ANOVA multiple comparisons and Kruskal-Wallis test and are indicated by asterisks: * p = < 0.05, ** p = < 0.01, **** p = < 0.0001. **C)** CLSTN mRNA expression analysis across the 12 MB subtypes. ANOVA multiple comparisons and Kruskal-Wallis test. * p = < 0.05, ** p = < 0.01, *** p = < 0.001, **** p = < 0.0001. **D)** XY plots showing the correlation between CLSTN and MAP4K4 expression across the four MB subgroups WNT, SHH, Grp3 and Grp4. Interrupted line indicates best fit of linear regression analysis. Slope and p-value of slope are indicated. **E)** Analysis of CLSTN1 expression across human organ development. **F, G)** snRNAseq analysis of CLSTN1 (E) and MAP4K4 (F) expression across cerebellar cell types. Upper: Manifold Approximation and Projection (UMAP) of 180,956 human cells colored by cell type (from^48^). Lower: Expression levels of CLSTN1 across projected cell types. CPM, Counts per million; GABA, Gamma-aminobutyric acid, DN, Dentate nucleus; GC, Granule cells; UBC, Unipolar brush cells; Glut_DN, Glutamatergic deep nuclei neurons; Isth_N, Isthmic nuclei neurons; NTZ, Nuclear transitory zone; VZ, Ventricular zone.

Taken together, this data indicates a reduced expression of CLSTN1 in MB tumor cells and points towards tight control of CLSTNs during cerebellar development. The negative correlation of *MAP4K4* and *CLSTN1* expression in the developing cerebellum further indicates a potential mechanism of mutual control of these two genes.

### CLSTN1 is associated with membranes in MB cells and detected at the cell surface

To explore expression levels of CLSTN1 in different established human MB tumor cell lines, we probed lysates of three SHH and three Grp3 MB cell lines with anti-CLSTN1 antibodies by immunoblotting (IB). We detected similar expression levels across all tested cell lines, with slightly higher expression in HD-MB03 (Fig. 2A, left). As we expected biological relevance from cell surface expressed CLSTN1 specifically, we also assessed CLSTN1 expression by flow cytometry (FC) analysis. We detected CLSTN1 expression on all six cell lines (Fig. 2B). Except for ONS-76, all cell lines displayed a small population of CLSTN1-low or -negative cells. The highest levels of CLSTN1 expression were detected in ONS-76 cells. To further confirm CLSTN1 membrane localization, we performed cell fractionation analysis in one SHH (UW228) and one Grp3 (HD-MB03) MB cell line. We detected CLSTN1 in the membrane fraction of both cell lines. Somewhat surprisingly, however, we detected a larger proportion of CLSTN1 in the cytosolic fraction in HD-MB03 (Fig. 2C). Immunofluorescence analysis revealed that a majority of CLSTN1 protein under regular growth conditions is localized diffusely in the cytoplasm with some enrichment in perinuclear regions (Fig. 2D). However, we also clearly detected CLSTN1 at the plasma membrane (arrowheads), in lamellipodia-like protrusions and filamentous cell-cell contacts (arrows).

**Figure 2:**
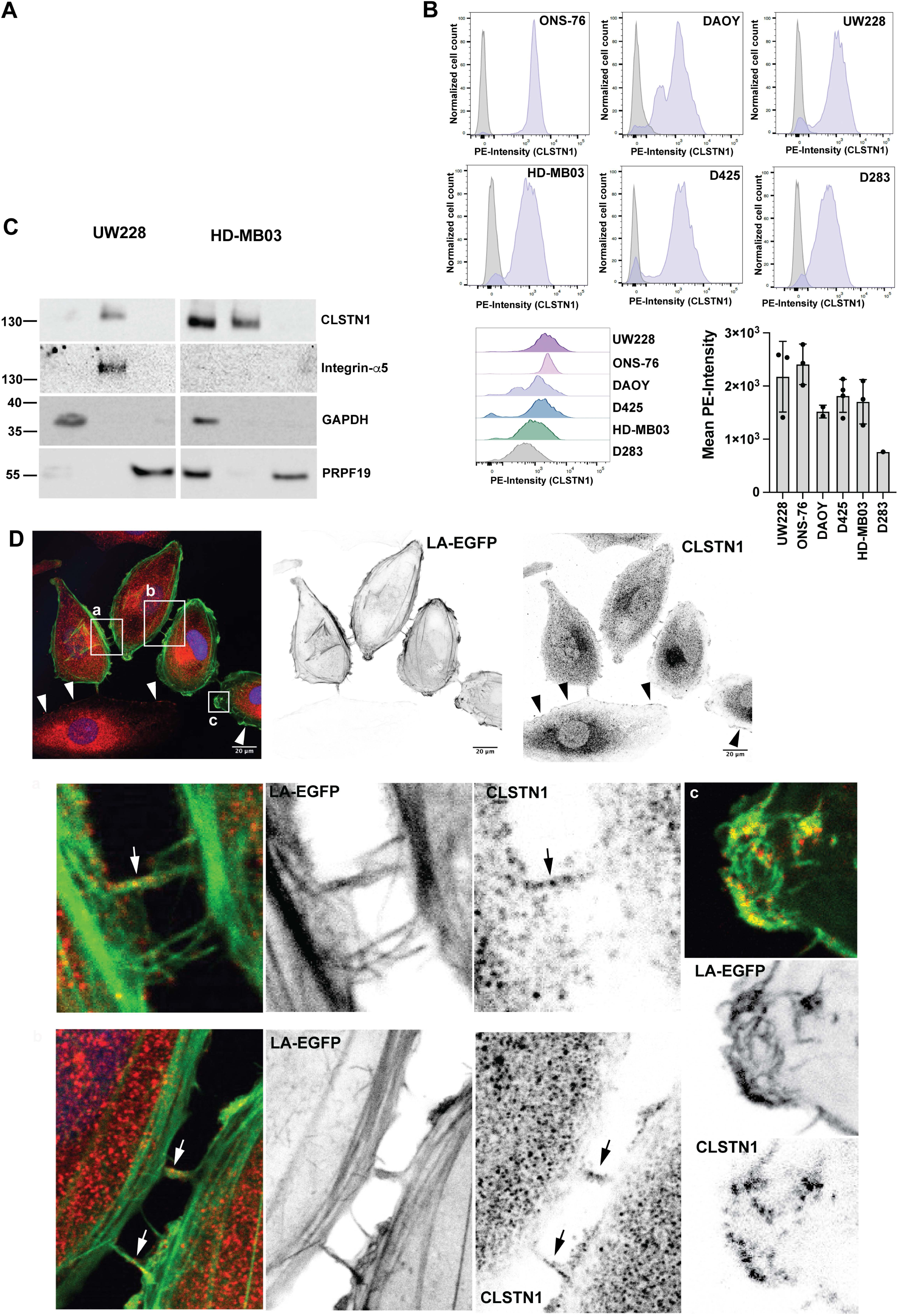
CLSTN1 accumulates at the cell surface, in internal small vesicle-like structures, in lamellipodia-like protrusions, and in filamentous cell-cell connections. **A)** Immunoblot (IB) analysis of CLSTN1 expression in six different MB cell lines. GAPDH expression was used as the loading control. **B)** Surface expression analysis of CLSTN1 in six different MB cell lines by FC. The bar plot depicts mean fluorescence intensities in the six different cell lines from different experiments. **C)** Subcellular fractionation analysis of CLSTN1 expression in the SHH MB cell line UW228 and the Grp3 MB cell line HD-MB03. Integrin-α5 and PRPF9 expression was used as fractionation controls for the membrane and the nuclear fractions, respectively. CEB: cytoplasmic extraction buffer; MEB: membrane extraction buffer; NEB: nuclear extraction buffer. **D)** IFA of endogenous CLSTN1 protein in UW228 LA-EGFP-mCherry-Nuc cells, where it accumulated in small vesicle-like, perinuclear structures. Inverted greyscale images of the green-fluorescent signal (LA-EGFP, middle) and the red-fluorescent signal (CLSTN1, right) are shown. Scale bar is 20 µm. Magnifications a, b and c depict CLSTN1 localization in filamentous cell-cell connections and lamellipodia-like protrusions, respectively. Arrowheads indicate CLSTN1 localization in the plasma membrane, arrows accumulations of CLSTN1 in the filamentous cell-cell connections.

### Reduced CLSTN1 expression increased invasion and reduced viability

To explore the potential functional significance of CLSTN1 in MB cells, we depleted CLSTN1 using two different siRNAs, which effectively reduced CLSTN1 protein in all tested cell lines (Fig. 3A). By flow cytometry, we could still detect CLSTN1 in siCLSTN1-treated cells. However the expression levels in siCLSTN1-transfected cells were significantly lower compared to control siRNA transfected cells (Fig. 3B). To assess the potential implication of CLSTN1 in cell invasion control, we compared the invasiveness of CLSTN1-depleted DAOY cells compared to control using the spheroid invasion assay (SIA). Interestingly, the reduction of CLSTN1 causes a significant increase in bFGF-induced invasion (Fig. 3C). Both si1-CLSTN1 and si2-CLSTN1 increased invasiveness to an extent, which correlates with the depletion efficacy (Fig. S3). As cells for the SIA are grown in low adhesion plates, we tested whether suspension culture could influence CLSTN1 protein levels. We found that cells grown as free-floating cell clusters expressed higher CLSTN1 protein levels (Fig. 3D). Depletion of CLSTN1 caused a moderate increase of focal connexin 43 (Cx43) staining in regions of cell-cell contact (Fig. S4A), but did not affect [Ca^2+^]_i_ fluctuations in the tumor cells (Fig. S4B). To assess the effect of CLSTN1 overexpression, we generated stable cell lines overexpressing either CLSTN1-mNeonGreen (CLSTN1-mNG) or CLSTN1-V5. A marked increase of CLSTN1 was detected in cells expressing CLSTN1-mNG (Fig. 3E). The band of the overexpressed CLSTN1-mNG was detected as expected at a higher molecular weight by IB (Fig. 3E, upper). The increased CLSTN1 protein expression caused a moderate increase in cell viability (Fig. 3F). Conversely, siRNA-mediated reduction of CLSTN1 reduced cell viability. To confirm that the overexpressed CLSTN1-mNG was localized similarly to the endogenous protein, we visualized CLSTN1-mNG localization in the cells by fluorescence microscopy (Fig. 3G). CLSTN1-mNG primarily accumulated around the nucleus. Comparable to the endogenous CLSTN1 (Fig. 2D), we also detected CLSTN1-mNG in the plasma membrane, in lamellipodia-like structures, and in cell-cell connections (Fig. 3G).

**Figure 3:**
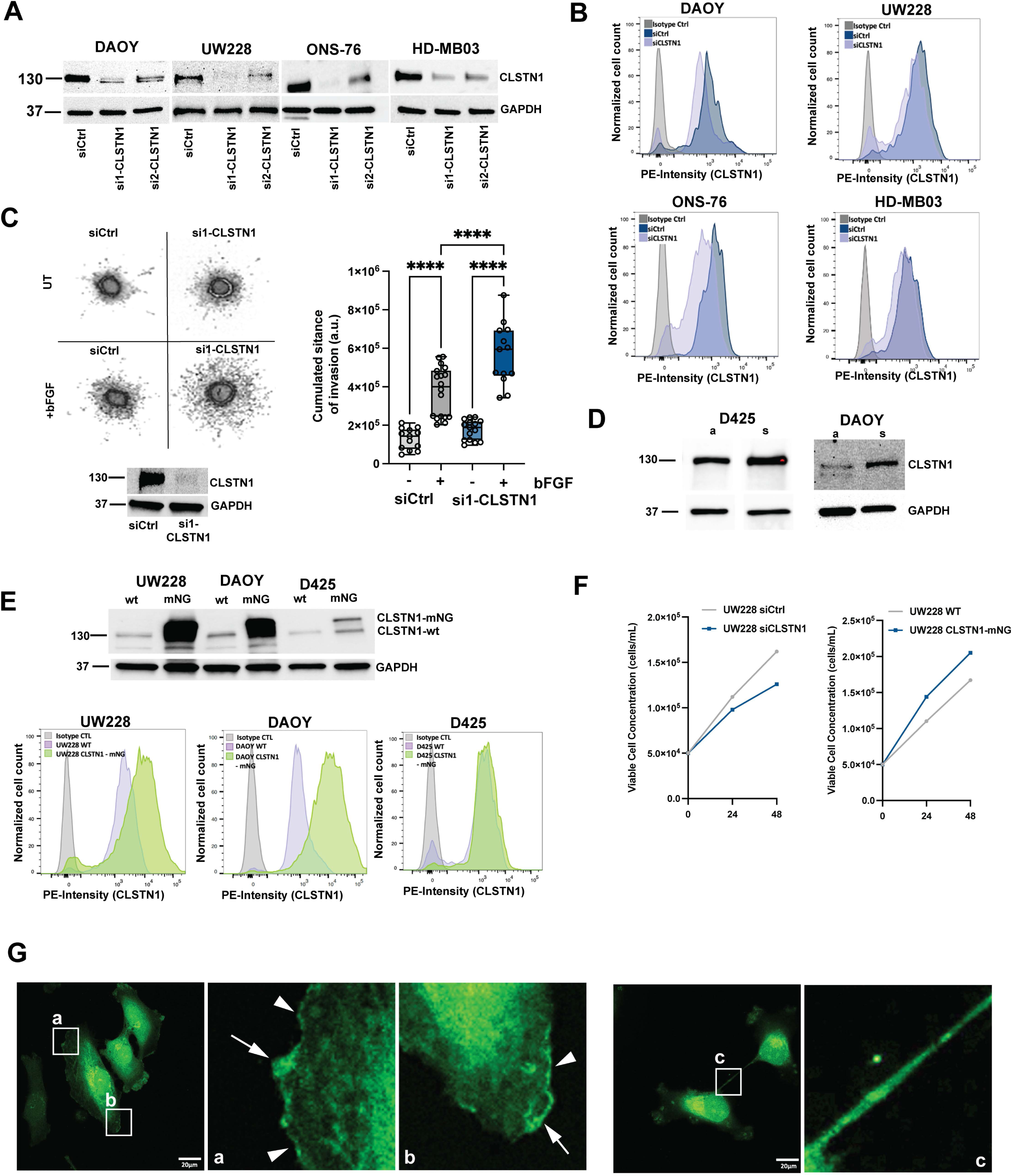
CLSTN1 expression levels determine the invasive phenotype of MB cells. IB analysis of CLSTN1 expression after siRNA-mediated depletion in DAOY, UW228, ONS-76 and HD-MB03 cell lines. Two different siRNAs targeting CLSTN1 were used. GAPDH expression was used as loading control. **B)** Surface expression analysis by FC of CLSTN1 after si-RNA-mediated depletion in DAOY, UW228, ONS-76, and HD-MB03 cell lines. **C)** SIA of bFGF stimulated (100 ng/ml) control (siCtrl) and si1-CLSTN1-transfected cells treated with 100 ng/ml bFGF for 24h. Upper left panel shows representative images of Hoechst-stained cultures at endpoint. UT: untreated. Lower panel is an IB validation of siRNA efficacy 48h after transfection. Right panel box-dot plot depicts the median cumulated invasion distances of n = 13 - 18 samples. ANOVA **** p = <0.0001. **D)** IB analysis of CLSTN1 expression in D425 and DAOY cells grown either as adherent or suspension cultures. E) IB (upper) and FC analysis (lower) IB analyses of endogenous (light purple) and ectopically overexpressed (light green) CLSTN1 in UW228, DAOY and D425 cells. **F)** Proliferation analysis of UW228 cells transfected with siCLSTN1 or of UW228 cells overexpressing CLSTN1-mNG. **G)** IFA of UW228 cells expressing CLSTN1-mNG. a, b and c depict magnifications of areas highlight with white rectangles. Arrowheads indicate CLSTN1-mNG localization at the plasma membrane, arrows localization of CLSTN1-mNG in lamellipodia-like structures. Magnification c depicts CLSTN1-mNG accumulation in filamentous cell-cell connection. Scale bar is 20 µm.

These data indicate that CLSTN1 is associated with cellular membranes and the plasma membrane. Depletion of CLSTN1 increases growth factor-induced invasion, suggesting that the expression levels of CLSTN1 may determine the invasive capabilities of the MB tumor cells.

### Visualization of CLSTN1 trafficking dynamics in living MB tumor cells

We used confocal live cell imaging of CLSTN1-mNG expressing UW228 cells to determine dynamic alterations in CLSTN1 localization in cells (Fig. 4A, movie 1). We detected CLSTN1-mNG accumulated in lamellipodia (a, movie 2), in a structure near the nucleus and reminiscent of the Golgi (b, movie 3), in small vesicles trafficking from the cell periphery towards the cell body (c, movie 4) and on filamentous structures, moving anterogradely relative to the nucleus (d, movie 5). Large endocytic vesicles in lamellipodia near the leading-edge membranes were devoid of fluorescence (movies 2 and 5), indicating that plasma membrane-localized CLSTN1 is internalized through a different endocytic mechanism. Rather, we observed markedly smaller, strongly fluorescent spots emanating from the lamellipodial membranes and trafficking inwards (Fig. 4A, movie 4). Fig. 4B and S5A depict time series of still images of an enlarged area (movie 7) from the original movie 6 (Fig. 4B). The filamentous, anterogradely moving structures are indicated by red arrowheads. Within the 10 min observation window, three consecutive passages of CLSTN1-mNG from the perinuclear region towards the cell periphery were observed (Fig. 4B).

**Figure 4:**
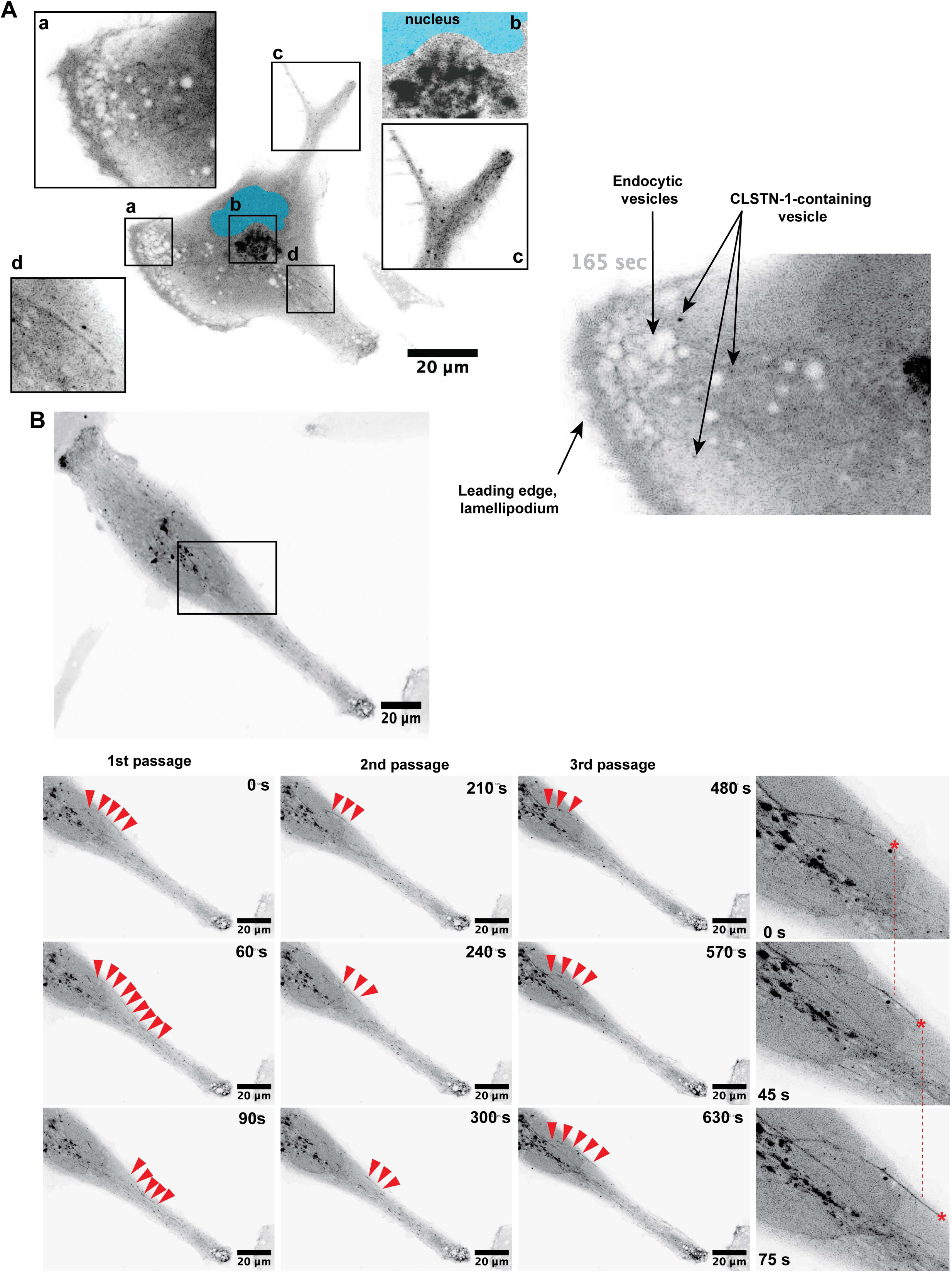
CLSTN1-mNG internalization and trafficking of CLSTN1-mNG. **A)** Still image of CLSTN1-mNG overexpressing UW228 cell. Inverted greyscale of green-fluorescent signal is shown. Squares a – d depict different subcellular localization of CLSTN1-mNG. a: Lamellipodia-like structure at cell leading edge; b: Perinuclear near Golgi; c: in small vesicles at cell trailing edge; d: on filamentous structures connecting cell core with periphery. B) Still image series of CLSTN1-mNG in UW228 cell. Inverted greyscale of green-fluorescent signal is shown. Red arrowheads indicate filamentous structure that is progressing from the perinuclear region toward cell periphery. Three passages of such structures are observed in 10 min observation period. Red asterisks indicate the position of front of filamentous structures at the indicated time point.

These observations point towards a rapid turnover of CLSTN1 at peripheral membranes. Delivery may occur through microtubule (+)-end-directed transport as was already suggested previously^14^, whereas internalization is likely mediated through a specific endocytic mechanism to be determined.

### CLSTN1 localizes to tumor-tumor cell and tumor-astrocytic cell contact zones

Using IFA, we next clarified the localization of endogenous CLSTN1 in tumor cells seeded as monocultures and tumor cells seeded in co-culture with immortalized normal human astrocytes (iNHA). In monoculture, most of the cell-cell contact zones displayed some CLSTN1 signal (Fig. 5A). However, this signal was rather weak and needed careful observation to be detected. In contrast, when cells were grown in co-culture with iNHA, the signal in tumor-tumor cell interaction zones increased both in intensity and abundance (Fig. 5B). We detected a CLSTN1 signal in nearly every cell-cell contact (Fig. 5B, S6A). In regions of tumor-iNHA cell interactions, we also detected CLSTN1 accumulations, indicating that CLSTN1 is localized to both homotypic and heterotypic cell interactions. We confirmed CLSTN1 localization to heterotypic tumor-astrocyte interactions using primary mouse cerebellar astrocytes (Fig. S6B). In homotypic cell-cell interactions between UW228 cells co-cultured with iNHA, CLSTN1 aligns in parallel linear structures of approximately 1 µm in length (Fig. 5B). By IB analysis we confirmed CLSTN1 expression in primary murine cerebellar astrocyte culture, which in expression levels is similar to the levels detected in the group 3 MB cell line HD-MB03 but considerably higher than in UW228 (Fig. 5C). In the cerebellar astrocyte culture, CLSTN1 migrates at a higher molecular weight in the SDS-PAGE, indicating either alternative splicing or differential posttranslational modifications.

**Figure 5:**
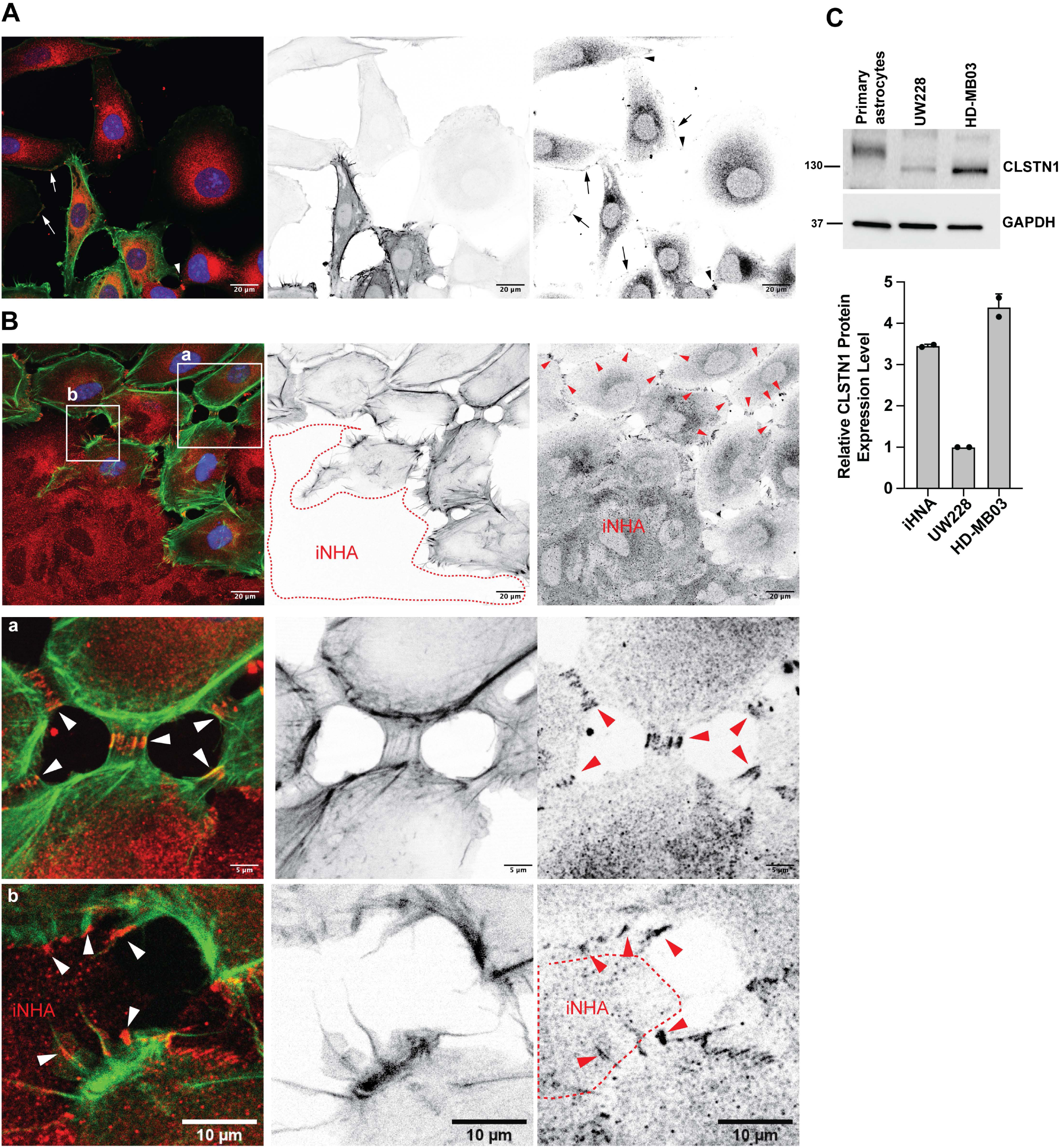
Increased CLSTN1 localization to cell-cell contacts in tumor-astrocyte co-cultures. **A)** IFA analysis of CLSTN1 expression in UW228-LA-EGFP-mCherry-Nuc cells. Red: CLSTNA1, green: actin (LA-EGFP), blue: Nuclei (mCherry-Nuc). Arrows indicate CLSTN1 localization at the cell cortex, arrowheads in regions of cell-cell contact. **B)** IFA analysis of CLSTN1 expression in UW228-LA-EGFP-mCherry-Nuc co-cultured with iNHA. iNHA are only positive for CLSTN1 and do not express LA-EGFP or mCherry-Nuc. Red dotted line indicates circumference of iNHA cell cluster, arrowheads indicate CLSTN1-positive cell-cell contacts. Magnifications of zones indicated with white rectangles show homotypic (a) and heterotypic (b) cell-cell interactions. **C)** IB analysis and quantification of CLSTn1 expression in primary murine cerebellar astrocytes, UW228 and Hd-MB03 MB cells.

Collectively, these data indicate that co-culture of UW228 tumor cells with iNHA increased CLSTN1 localization to cell-cell contacts.

### Up-regulation of CLSTN1 in cell-cell contacts upon prostetin/12k treatment

Surface proteomic analysis in DAOY MB cells indicated that CLSTN1 surface expression is negatively regulated by the Ser/Thr kinase MAP4K4^30^. MAP4K4 is a pro-invasive kinase with pleiotropic functions in health and disease^31^, highly expressed in MB, where it drives growth factor-induced cell invasion^32,33^. To test whether MAP4K4 kinase activity could be involved in the control of CLSTN1 plasma membrane expression, we treated cells with a novel pan MAP4K inhibitor prostetin/12K^34^ for 24 h and determined CLSTN1 expression and subcellular distribution by cell fractionation analysis (Fig. 6A). CLSTN1 was detectable exclusively in the membrane fraction in UW228 cells, where we observed a moderate increase in cells treated with prostetin/12K. To confirm that increased CLSTN1 in the membrane fraction translates into increased cell surface expression, we probed prostetin/12K-treated cells by FC analysis. We found that prostetin/12K treatment shifted the relative mean fluorescence intensity of CLSTN1 in UW228 by 46% (n = 3 independent experiments, SD = 7.3%, Fig. 6B,C) and by 47% in DAOY cells (Fig. S5B). siRNA-mediated depletion of MAP4K4 also causes a small increase in CLSTN1 surface expression by 23% (n = 3 independent experiments, SD = 4,.2%, Fig. S5C,D). We next compared prostetin/12K impact on subcellular CLSTN1 distribution by IFA. In DMSO-treated samples, we detected CLSTN1 in regions of cell-cell contact (Fig. 6D). Prostetin/12K treatment caused a marked increase in cell size and led to the accumulation of CLSTN1-mNG in regions of cell-cell contact (Fig. 6E, S6C). CLSTN1 accumulation was particularly evident in contacts between tumor cells and iNHA after prostetin/12K treatment (Fig. 6F) where it focally accumulated specifically in contacts between UW228 and iNHA (Fig. S7A,B). Increased CLSTN1 expression is not a consequence of increased *CLSTN1* transcription as we did not observe a significant increase in CLSTN1 mRNA after 6 and 24 h prostetin/12K treatment (Fig. S5E).

**Figure 6:**
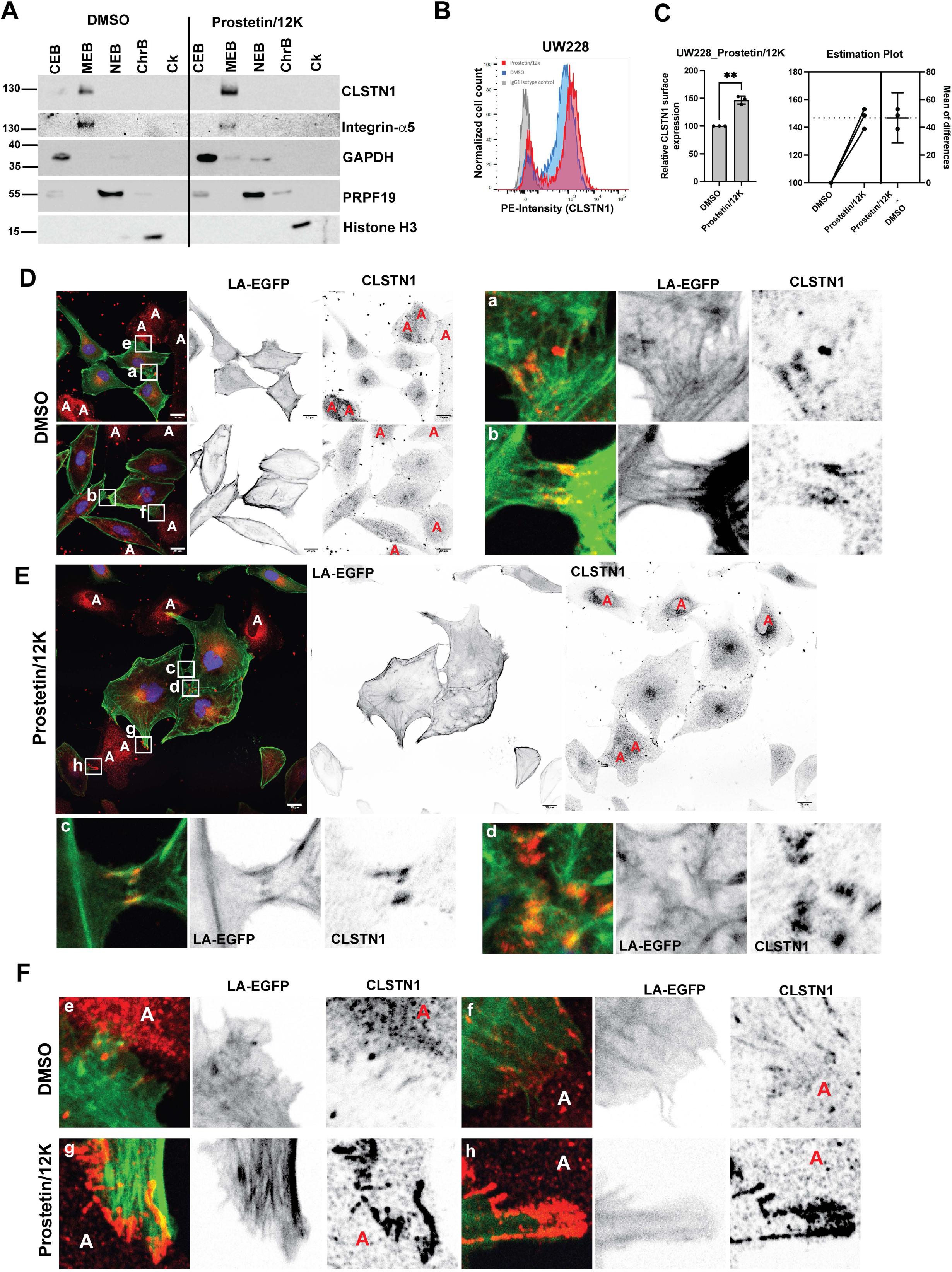
Pharmacological or genetic perturbation of MAP4K4 increased CLSTN1 expression and cell-cell contact accumulation. **A)** Cell fractionation analysis of CLSTN1 subcellular localization in UW228 cells. Antibodies against proteins indicated to the right of the panels were used. **B)** FC analysis of CLSTN1 surface expression on UW228 cells treated with 0.5 µM prostetin/12K for 24h. **C)** Quantification of relative CLSTN1 surface expression from FC analysis shown in B and estimation plot of n = 3 independent experiments. **D)** IFA analysis of CLSTN1 expression in DMSO-treated (24h) UW228-LA-EGFP-mCherry-Nuc cells co-cultured with iNHA. Red: CLSTNA1, green: actin (LA-EGFP), blue: Nuclei (mCherry-Nuc). Inverted greyscale images of the green-fluorescent signal (LA-EGFP, middle) and the red-fluorescent signal (CLSTN1, right) are shown. The scale bar is 20 µm. Yellow asterisks indicate the positions of iNHAs. Magnifications in a) and b) indicate cell-cell contacts between tumor cells. **E)** FA analysis of CLSTN1 expression in 0.5 µM prostetin/12K-treated (24h) UW228-LA-EGFP-mCherry-Nuc cells co-cultured with iNHA. Red: CLSTNA1, green: actin (LA-EGFP), blue: Nuclei (mCherry-Nuc). Yellow asterisks indicate the positions of iNHAs. Magnifications in c) and d) indicate cell-cell contacts between tumor cells. **F)** Magnifications of tumor-iNHA cell-cell contacts in DMSO- (e and f) or prostetin/12K- (g and h) treated samples. Yellow asterisks indicate the positions of iNHAs.

Collectively these data indicate that the prostetin/12K target MAP4K4 could be involved in CLTSN1 expression and function in MB and control subcellular localization of CLSTN1 in cell-cell contacts.

## Discussion

Localization, regulation, and functions of the neuronal postsynaptic adhesion protein CLSTN1 have been mostly investigated in the context of physiological cell functions. Herein we addressed the expression, subcellular localization, and function of CLSTN1 in the embryonal brain tumor medulloblastoma and further explored its previously observed regulation by the pro-invasive kinase MAP4K4. We provide evidence that CLSTN1 may exert tumor-suppressive functions by repressing invasiveness and enabling homo- and heterotypic cell-cell interactions. These functions of CLSTN1 are in part controlled by MAP4 kinases, as prostetin/12K, a neuroprotective, novel selective inhibitor of this kinase family, increased plasma membrane-associated CLSTN1 expression in MB tumor cells and enhanced CLSTN1 localization in tumor-astrocyte interactions. Depletion of CLSTN1 on the other hand caused a significant increase in growth factor-induced invasion. Together, our study provides novel insights into the functions of CLSTN1 in the context of a malignant brain tumor. Specifically, it sheds light on kinase regulation of CLSTN1 subcellular abundance and localization, which we found to be of potential importance in mediating microenvironment interactions.

In the differentiating chick cerebellum, CLSTN1 was found to be strongly expressed in granule cells in the internal granule layer (IGC), whereas CLSTN2 and CLSTN3 are specifically expressed in Purkinje cells^35^. This finding was recently confirmed in the human cerebellum, where CLSTN1 is predominately expressed in a subset of granule cells (GCs)^29^. In contrast, CLSTN2 was found localized to a subset of Purkinje cells, unipolar brush cells, and a subset of granule cells. CLSTN3 was predominately detected in a subset of granule cells and dispersed in some Purkinje cell populations. High CLSTN1 expression in cerebellar GCs is contrasted by low MAP4K4 expression in the same cellular compartment^29^, which corroborates the negative regulatory function of MAP4K4 on CLSTN1 expression we observed *in vitro*. The overall negative correlation of MAP4K4 and CLSTN1 expression *in vivo* and our findings of increased CLSTN1 protein levels after prostetin/12K treatment indicate that CLSTN1 expression could be negatively regulated by MAP4 kinases. We observed no increase in CLSTN1 transcripts after 6 or 24h of prostetin/12K treatment, suggesting that CLSTN1 protein expression is controlled by MAP4K via a posttranscriptional mechanism. A study in whole mice and lung adenocarcinoma cells found that CLSTN1 expression is decreased in response to 4 Gy gamma irradiation^36^. We previously observed an increased MAP4K4-dependent invasiveness in SHH MB tumor cells exposed to 1 – 2 Gy^37^, suggesting that the irradiation-induced increase in cell invasion observed could in part be mediated by CLSTN1 downregulation via increased MAP4K4 activity.

Our live-cell imaging analysis of the subcellular dynamics of CLSTN1 confirms anterograde trafficking in MB tumor cells. This observation could indicate that CLSTN1 links organelles to kinesin-mediated transport processes and cargo delivery in tumor cells as was observed in healthy neuronal axons^12,13^. Consequently, the increased accumulation of CLSTN1 in homo- and heterotypic cell-cell interactions upon prostetin/12K treatment may indicate that MAP4Ks either prevent proper trafficking of CLSTN1 and associated cargo or cause increased turnover at adhesion sites by controlling proteins involved in CLSTN1 trafficking, such as ERK, which negatively regulates CLSTN1-KLC via KLC1S460 phosphorylation^15^. The repression of ERK was shown to reduce S460 phosphorylation and increase CLSTN1-KLC1 binding, suggesting that CLSTN1 trafficking could be controlled by reversible KLC1 S460 phosphorylation downstream of MAPK signaling^15^. MAP4K4 activates ERK through repression of PP2A in lung adenocarcinoma^38^; hence, prostetin/12K treatment may increase CLSTN1 interaction with KCL1 through the reduction of S460 phosphorylation in KLC1.

We observed a small increase in the gap junction protein Cx43 in cell-cell contact zones after CLSTN1 depletion. As we observed no changes in [Ca^2+^]_i_, we do not think that CLSTN1 is implicated in [Ca^2+^]_i_ propagation in MB tumor cells. Thus, the functional significance of CLSTN1 recruitment and accumulation in regions of cell-cell contacts between tumor cells and astrocytes remains to be determined. The high number of astrocytes in the cerebellum, which outnumber Purkinje cells and provide critical functions that support the proliferation and migration of cerebellar granule neuron precursors^39^, argues for the relevance of direct tumor-astrocyte interaction in the control of tumor growth and progression. Indeed, in MB with extensive nodularity (MBEN), astrocytic cells were found associated with malignant and migratory cells in the internodular space, which was suggested to provide a source of neoplastic cells that then migrate to the nodular compartment while losing the proliferative potential^40^. Additionally, MB tumor-associated astrocytes display even distribution in small SHH tumors or accumulation at the tumor margin in large tumors, where they form a glial scar in mouse models *in vivo*^41^. Importantly, several studies identified SHH MB tumor-promoting activities of astrocytes via the induction of sustained hedgehog signaling^41–44^. Loss of Patched is sufficient for the induction of SHH medulloblastoma from granule neuronal precursor cells^45^, but astrocyte secretion of SHH is necessary to support MB growth, and astrocyte ablation prevented progression of SHH tumors in Ptch1^−/−^ mice^46^. Astrocyte-secreted fibronectin supports the outgrowth and survival of primary SHH-derived tumoroids^47^, suggesting that both soluble and physical components secreted by astrocytes contribute to MB tumor growth. Future studies should thus address how the tumor-astrocyte coupling via CLSNT1 in cells treated with the MAP4K inhibitor prostetin/12K could influence hedgehog signaling as well as astrocyte biology and function in the context of MB.

In conclusion, our study provides a first insight into potential tumor-restrictive interactions between tumor cells and astrocytes, which involve CLSTN1 and its regulation by the MAP4 kinase family.

## Figure legends

**Figure S1: CLSTN2 and CLSTN3 are differentially expressed during organ development.** Analysis of CLSTN2 **(A)** and CLSTN3 **B)** expression across human organ development. snRNAseq analysis of CLSTN2 **(C)** and CLSTN3 **(D)** expression across cerebellar cell types expression. Left: Manifold Approximation and Projection (UMAP) of 180,956 human cells colored by cell type (from^48^). Right: Expression levels of CLSTN1 across projected cell types. CPM, Counts per million; GABA, Gamma-aminobutyric acid, DN, Dentate nucleus; GC, Granule cells; UBC, Unipolar brush cells; Glut_DN, Glutamatergic deep nuclei neurons; Isth_N, Isthmic nuclei neurons; NTZ, Nuclear transitoryzone; VZ, Ventricular zone.

**Figure S2:** Pseudobulk gene expression analysis of CLSTN1, CLSTN2 and CLSTN3 along cerebellar development.

**Figure S3: si1-CLSTN1 and si2-CLSTN1 are comparably efficient in increasing bFGF-induced invasion of the tumor cells.** Upper panel: SIA of DAOY cells stimulated with 100 ng/ml bFGF for 24 h. Representative images of Hoechst-stained cells at endpoint are shown. Lower left panel depicts the IB validation of siRNA efficacy 48h after transfection. Lower right panel box-dot plot depicts median cumulated invasion distances of n = 7 - 10 samples. ANOVA **** p = <0.0001.

**Figure S4: Depletion of CLSTN1 causes a moderate increase in CX43 in cell-cell contacts without affecting Ca^2+^ signaling. A)** IFA analysis of CX43 expression in siCTL or si1_CLSTN1-transfected UW228 cells. Green: LA-EGFP, cyan: CX43. Yellow arrowheads point towards CX43-positive cell-cell contacts. **B)** Quantification of [Ca^2+^]_i_ using the genetically encoded GCaMP6 calcium sensor in UW228 cells.

**Figure S5: CLSTN1-mNG trafficking on filamentous structures towards cell leading edge. A)** Still image of CLSTN1-mNG overexpressing UW228 cell. Inverted grey-scale of green-fluorescent signal is shown. Red arrowheads point to the front of moving CLSTN1-mNG on the filamentous structure. **B)** FC analysis of CLSTN1 surface expression on DAOY cells treated with 0.5 µM prostetin/12K for 24h. **C)** FC analysis of CLSTN1 surface expression on UW228 cells transfected either with siCTRL or siMAP4K4. **D)** Quantification of relative CLSTN1 surface expression from FC analysis shown in B and estimation plot of n = 3 independent experiments. **E)** qRT-PCR analysis of CLSTN1 mRNA expression after 6 and 24h of prostetin/12K treatment. Mean and SD of N = 3 technical replicas.

**Figure S6: A)** IFA analysis of CLSTN1 expression in UW228-LA-EGFP-mCherry-Nuc cells co-cultured with iNHA. Red: CLSTNA1, green: actin (LA-EGFP), blue: Nuclei (mCherry-Nuc). Letter “A “indicates the positions of iNHAs. Magnification in a) highlights cell-cell contacts between tumor cells. Arrowheads point towards CLSTN1-positive cell-cell contacts between tumor cells. **B)** IFA analysis of CLSTN1 expression in UW228-LA-EGFP-mCherry-Nuc cells co-cultured with primary murine cerebellar astrocytes. Red: CLSTNA1, green: actin (LA-EGFP), blue: Nuclei (mCherry-Nuc), purple: GFAP. Square highlights CLSTN1-positive contact between UW228 tumor cells and astrocytes. C) Comparative IFA analysis of CLSTN1 expression in UW228-LA-EGFP-mCherry-Nuc cells co-cultured with iNHA, without or with 0.5 µM prostetin/12K for 24h. Red: CLSTNA1, green: actin (LA-EGFP), blue: Nuclei (mCherry-Nuc). Arrows point towards CLSTN1-positive cell-cell contacts between UW228 and iNHAs. Letter “A “indicates the positions of iNHAs.

**Figure S7: A)** and **B)** show two different samples of UW228-LA-EGFP-mCherry-Nuc cells co-cultured with iNHA in the presence of 0.5 µM prostetin/12K for 24h. Magnifications a) UW228-iNHA CLSTN1-positive cell-cell contacts, magnifications b) UW228-UW228 CLSTN1-positive cell-cell contacts. Letter “A “indicates the positions of iNHAs.

## References

1. Cavalli, F. M. G. et al. Intertumoral Heterogeneity within Medulloblastoma Subgroups. Cancer Cell 31, 737–754.e6 (2017).

2. Fults, D. W., Taylor, M. D. & Garzia, L. Leptomeningeal dissemination: A sinister pattern of medulloblastoma growth. J Neurosurg Pediatr 23, 613–621 (2019).

3. Vladoiu, M. C. et al. Childhood cerebellar tumours mirror conserved fetal transcriptional programs. Nature 572, 67–73 (2019).

4. Hovestadt, V. et al. Resolving medulloblastoma cellular architecture by single-cell genomics. Nature 572, 74–79 (2019).

5. Zhang, L. et al. Single-Cell Transcriptomics in Medulloblastoma Reveals Tumor-Initiating Progenitors and Oncogenic Cascades during Tumorigenesis and Relapse Article Single-Cell Transcriptomics in Medulloblastoma Reveals Tumor-Initiating Progenitors and Oncogenic Cascades. Cancer Cell 36, 302–318.e7 (2019).

6. Ocasio, J. et al. scRNA-seq in medulloblastoma shows cellular heterogeneity and lineage expansion support resistance to SHH inhibitor therapy. Nat Commun 10, 1–17 (2019).

7. Gojo, J. et al. Single-Cell RNA-Seq Reveals Cellular Hierarchies and Impaired Developmental Trajectories in Pediatric Ependymoma. Cancer Cell 38, 44–59.e9 (2020).

8. Jessa, S., et al. Stalled Developmental Programs at the Root of Pediatric Brain Tumors. Nature Genetics vol. 51 (Springer US, 2019).

9. Capdeville, C. et al. Spatial proteomics finds CD155 and Endophilin-A1 as mediators of growth and invasion in medulloblastoma. Life Sci Alliance 5, e202201380 (2022).

10. Vogt, L. et al. Calsyntenin-1, a proteolytically processed postsynaptic membrane protein with a cytoplasmic calcium-binding domain. Molecular and Cellular Neuroscience 17, 151–166 (2001).

11. Hintsch, G. et al. The calsyntenins - A family of postsynaptic membrane proteins with distinct neuronal expression patterns. Molecular and Cellular Neuroscience 21, 393–409 (2002).

12. Filigheddu, N. et al. Calsyntenin-1 Docks Vesicular Cargo to Kinesin-1. Mol Biol Cell 18, 986– 994 (2007).

13. Araki, Y. et al. The novel cargo Alcadein induces vesicle association of kinesin-1 motor components and activates axonal transport. EMBO Journal 26, 1475–1486 (2007).

14. Ludwig, A. et al. Calsyntenins mediate TGN exit of APP in a kinesin-1-dependent manner. Traffic 10, 572–589 (2009).

15. Vagnoni, A., Rodriguez, L., Manser, C., De Vos, K. J. & Miller, C. C. J. Phosphorylation of kinesin light chain 1 at serine 460 modulates binding and trafficking of calsyntenin-1. J Cell Sci 124, 1032–1042 (2011).

16. Mórotz, G. M. et al. Kinesin light chain-1 serine-460 phosphorylation is altered in Alzheimer’s disease and regulates axonal transport and processing of the amyloid precursor protein. Acta Neuropathol Commun 7, 200 (2019).

17. Ster, J. et al. Calsyntenin-1 regulates targeting of dendritic NMDA receptors and dendritic spine maturation in CA1 hippocampal pyramidal cells during postnatal development. Journal of Neuroscience 34, 8716–8727 (2014).

18. Ponomareva, O. Y., Holmen, I. C., Sperry, A. J., Eliceiri, K. W. & Halloran, M. C. Calsyntenin-1 regulates axon branching and endosomal trafficking during sensory neuron development in vivo. Journal of Neuroscience 34, 9235–9248 (2014).

19. Cheng, K. et al. Calsyntenin-1 Negatively Regulates ICAM5 Accumulation in Postsynaptic Membrane and Influences Dendritic Spine Maturation in a Mouse Model of Fragile X Syndrome. Front Neurosci 13, 1–16 (2019).

20. Li, C. et al. ESRP1-driven alternative splicing of CLSTN1 inhibits the metastasis of gastric cancer. Cell Death Discov 9, (2023).

21. Hu, X. et al. The RNA-binding protein AKAP8 suppresses tumor metastasis by antagonizing EMT-associated alternative splicing. Nat Commun 11, (2020).

22. Tripolitsioti, D. et al. MAP4K4 controlled integrin β1 activation and c-Met endocytosis are associated with invasive behavior of medulloblastoma cells. Oncotarget 9, 23220–23236 (2018).

23. Migliavacca, J., Züllig, B., Capdeville, C., Grotzer, M. A. & Baumgartner, M. Cooperation of Striatin 3 and MAP4K4 promotes growth and tissue invasion. Commun Biol 5, (2022).

24. Kumar, K. S. et al. Computer-assisted quantification of motile and invasive capabilities of cancer cells. Sci Rep 5, (2015).

25. Sonoda, Y. et al. Formation of intracranial tumors by genetically modified human astrocytes defines four pathways critical in the development of human anaplastic astrocytoma. Cancer Res 61, 4956–4960 (2001).

26. Cavalli, F. M. G. et al. Intertumoral Heterogeneity within Medulloblastoma Subgroups. Cancer Cell 31, 737--754.e6 (2017).

27. Cardoso-Moreira, M. et al. Gene expression across mammalian organ development. Nature 571, 505–509 (2019).

28. Jovanovic, D., Yan, S. & Baumgartner, M. The molecular basis of the dichotomous functionality of MAP4K4 in proliferation and cell motility control in cancer. Frontiers in Oncology vol. 12 1–18 Preprint at 10.3389/fonc.2022.1059513 (2022).

29. Sepp, M. et al. Cellular development and evolution of the mammalian cerebellum. Nature (2023) doi:10.1038/s41586-023-06884-x.

30. Capdeville, C. et al. Spatial proteomics finds CD155 and Endophilin-A1 as mediators of growth and invasion in medulloblastoma. Life Sci Alliance 5, e202201380 (2022).

31. Jovanovic, D., Yan, S. & Baumgartner, M. The molecular basis of the dichotomous functionality of MAP4K4 in proliferation and cell motility control in cancer. Frontiers in Oncology vol. 12 1–18 Preprint at 10.3389/fonc.2022.1059513 (2022).

32. 32. Migliavacca, J., Züllig, B., Capdeville, C., Grotzer, M. A. & Baumgartner, M. Cooperation of Striatin 3 and MAP4K4 promotes growth and tissue invasion. Commun Biol 5, (2022).

33. Tripolitsioti, D. et al. MAP4K4 controlled integrin β1 activation and c-Met endocytosis are associated with invasive behavior of medulloblastoma cells. Oncotarget 9, 23220–23236 (2018).

34. Bos, P. H. et al. Development of MAP4 Kinase Inhibitors as Motor Neuron-Protecting Agents. Cell Chem Biol 26, 1703–1715.e37 (2019).

35. De Ramon Francàs, G., Alther, T. & Stoeckli, E. T. Calsyntenins are expressed in a dynamic and partially overlapping manner during neural development. Front Neuroanat 11, 1–13 (2017).

36. Lee, E.-S., Kim, W.-T., Park, G.-Y., Lee, M. & Gen Son, T. Calsyntenin 1 mRNA expression sensitivity to ionizing radiation in human hepatocytes and carcinoma cells and blood cells of BALB/c mice. J Radiat Res Appl Sci 14, 44–50 (2021).

37. Tripolitsioti, D. et al. MAP4K4 controlled integrin β1 activation and c-Met endocytosis are associated with invasive behavior of medulloblastoma cells. Oncotarget 9, 23220–23236 (2018).

38. Gao, X. et al. MAP4K4 is a novel MAPK/ERK pathway regulator required for lung adenocarcinoma maintenance. Mol Oncol 11, 628–639 (2017).

39. De Zeeuw, C. I. & Hoogland, T. M. Reappraisal of Bergmann glial cells as modulators of cerebellar circuit function. Frontiers in Cellular Neuroscience vol. 9 1–8 Preprint at 10.3389/fncel.2015.00246 (2015).

40. Ghasemi, D. R. et al. Compartments in medulloblastoma with extensive nodularity are connected through differentiation along the granular precursor lineage. Nat Commun 15, 1–20 (2024).

41. Li, H. et al. Tumor-associated astrocytes promote tumor progression of Sonic Hedgehog medulloblastoma by secreting lipocalin-2. Brain Pathology 1–19 (2023) doi:10.1111/bpa.13212.

42. Guo, D. et al. Tumor cells generate astrocyte-like cells that contribute to SHH-driven medulloblastoma relapse. Journal of Experimental Medicine 218, (2021).

43. Cheng, Y. et al. Sustained hedgehog signaling in medulloblastoma tumoroids is attributed to stromal astrocytes and astrocyte-derived extracellular matrix. Laboratory Investigation 100, 1208–1222 (2020).

44. Liu, Y. et al. Astrocytes Promote Medulloblastoma Progression through Hedgehog Secretion. Cancer Res 77, 6692–6703 (2017).

45. Yang, Z. J. et al. Medulloblastoma Can Be Initiated by Deletion of Patched in Lineage-Restricted Progenitors or Stem Cells. Cancer Cell 14, 135–145 (2008).

46. Liu, Y. et al. Astrocytes Promote Medulloblastoma Progression through Hedgehog Secretion. Cancer Res 77, 6692–6703 (2017).

47. Cheng, Y. et al. Sustained hedgehog signaling in medulloblastoma tumoroids is attributed to stromal astrocytes and astrocyte-derived extracellular matrix. Laboratory Investigation (2020) doi:10.1038/s41374-020-0443-2.

48. Sepp, M. et al. Cellular development and evolution of the mammalian cerebellum. Nature 1–49 (2023) 10.1038/s41586-023-06884-x.

