## Supplementary figures for "MAP4 kinase-regulated reduced CLSTN1 expression in medulloblastoma is associated with increased invasiveness"

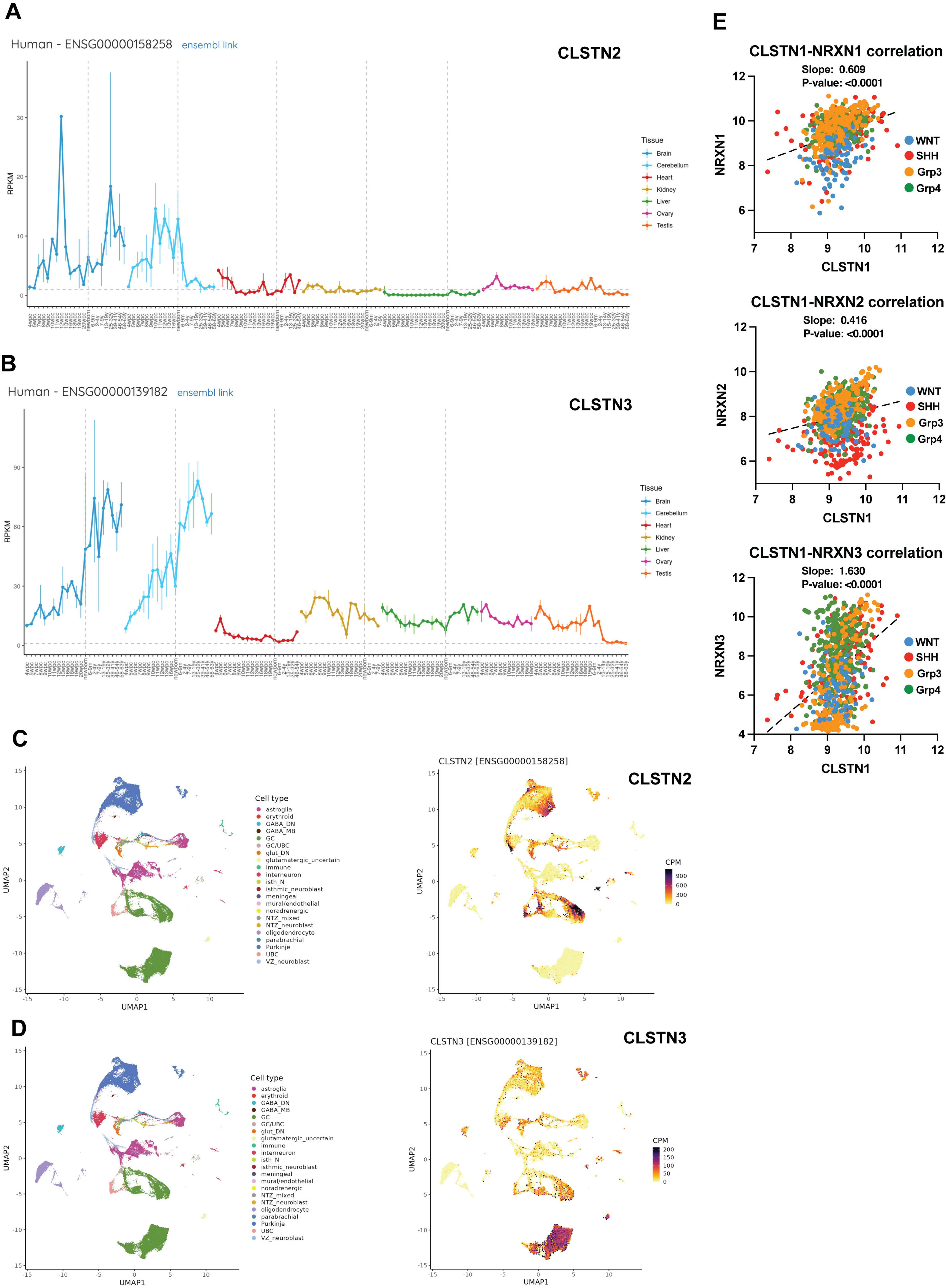
A
Human - ENSG00000158258 ensembl link
CLSTN2
RPKM
Tissue
Brain
Cerebellum
Heart
Kidney
Liver
Ovary
Testis
B
Human - ENSG00000139182 ensembl link
CLSTN3
RPKM
Tissue
Brain
Cerebellum
Heart
Kidney
Liver
Ovary
Testis
C
Cell type
astroglia
erythroid
GABA_DN
GABA_MB
GC
GC/UBC
glut_DN
glutamatergic_uncertain
immune
interneuron
isth_N
isthmic_neuroblast
meningeal
mural/endothelial
noradrenergic
NTZ_mixed
NTZ_neuroblast
oligodendrocyte
parabrachial
Purkinje
UBC
VZ_neuroblast
UMAP1
UMAP2
CLSTN2 [ENSG00000158258]
CLSTN2
CPM
D
Cell type
UMAP1
UMAP2
CLSTN3 [ENSG00000139182]
CLSTN3
CPM
E
CLSTN1-NRXN1 correlation
Slope: 0.609
P-value: <0.0001
NRXN1
CLSTN1
WNT
SHH
Grp3
Grp4
CLSTN1-NRXN2 correlation
Slope: 0.416
P-value: <0.0001
NRXN2
CLSTN1
WNT
SHH
Grp3
Grp4
CLSTN1-NRXN3 correlation
Slope: 1.630
P-value: <0.0001
NRXN3
CLSTN1
WNT
SHH
Grp3
Grp4

Fig. S1

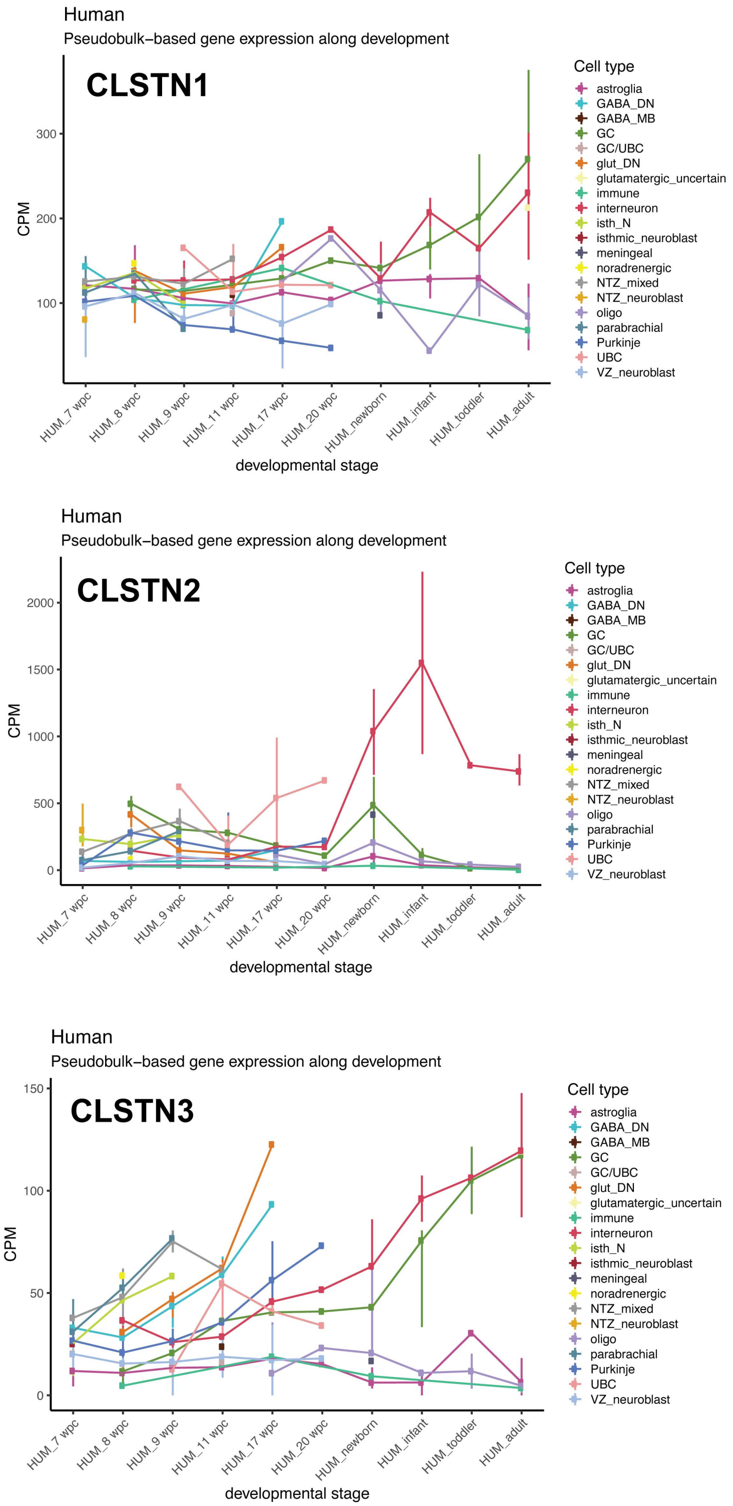
Human
Pseudobulk−based gene expression along development
CLSTN1
CPM
100
200
300
HUM_7 wpc
HUM_8 wpc
HUM_9 wpc
HUM_11 wpc
HUM_17 wpc
HUM_20 wpc
HUM_newborn
HUM_infant
HUM_toddler
HUM_adult
developmental stage
Cell type
astroglia
GABA_DN
GABA_MB
GC
GC/UBC
glut_DN
glutamatergic_uncertain
immune
interneuron
isth_N
isthmic_neuroblast
meningeal
noradrenergic
NTZ_mixed
NTZ_neuroblast
oligo
parabrachial
Purkinje
UBC
VZ_neuroblast
Human
Pseudobulk−based gene expression along development
CLSTN2
CPM
0
500
1000
1500
2000
developmental stage
Human
Pseudobulk−based gene expression along development
CLSTN3
CPM
0
50
100
150
developmental stage

Fig. S2

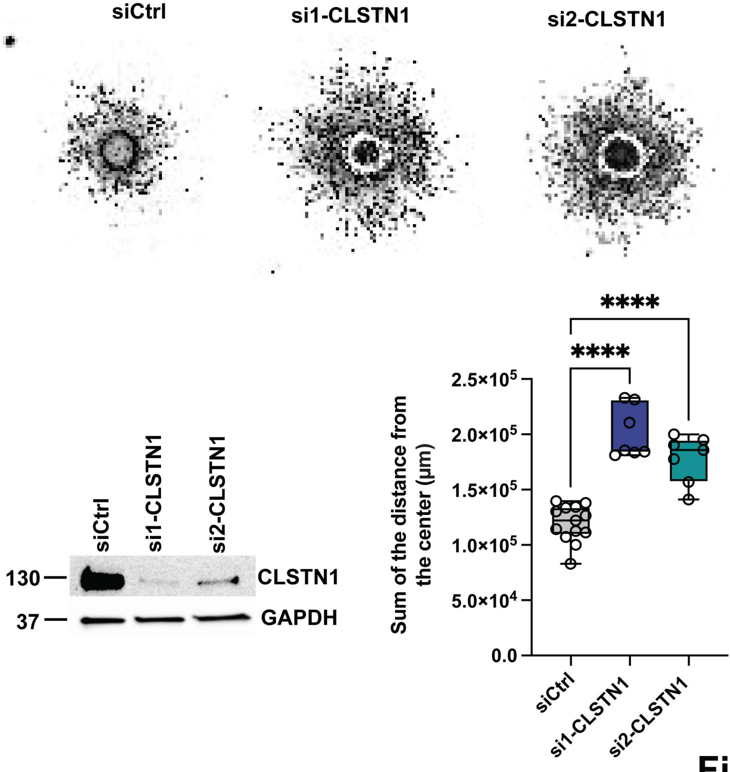
siCtrl
si1-CLSTN1
si2-CLSTN1
siCtrl
si1-CLSTN1
si2-CLSTN1
130
37
CLSTN1
GAPDH
****
****
Sum of the distance from the center (μm)
2.5×10^5
2.0×10^5
1.5×10^5
1.0×10^5
5.0×10^4
0.0
siCtrl
si1-CLSTN1
si2-CLSTN1

**Fig. S3**

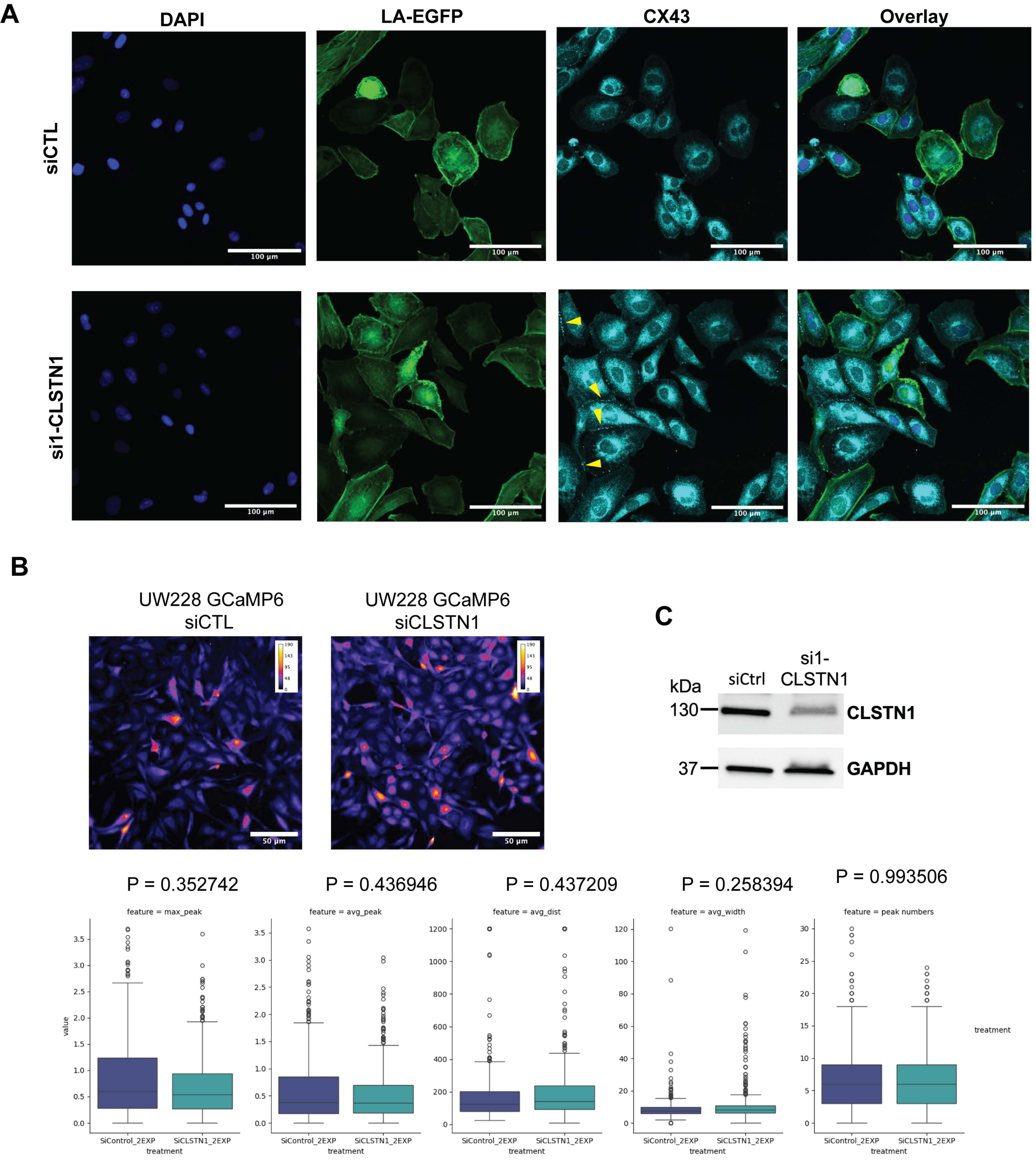
A
DAPI
LA-EGFP
CX43
Overlay
siCTL
si1-CLSTN1
100 µm
B
UW228 GCaMP6 siCTL
UW228 GCaMP6 siCLSTN1
50 µm
C
kDa
siCtrl
si1-CLSTN1
130
37
CLSTN1
GAPDH
P = 0.352742
P = 0.436946
P = 0.437209
P = 0.258394
P = 0.993506
feature = max_peak
feature = avg_peak
feature = avg_dist
feature = avg_width
feature = peak numbers
value
treatment
SiControl_2EXP
SiCLSTN1_2EXP

Fig. S4

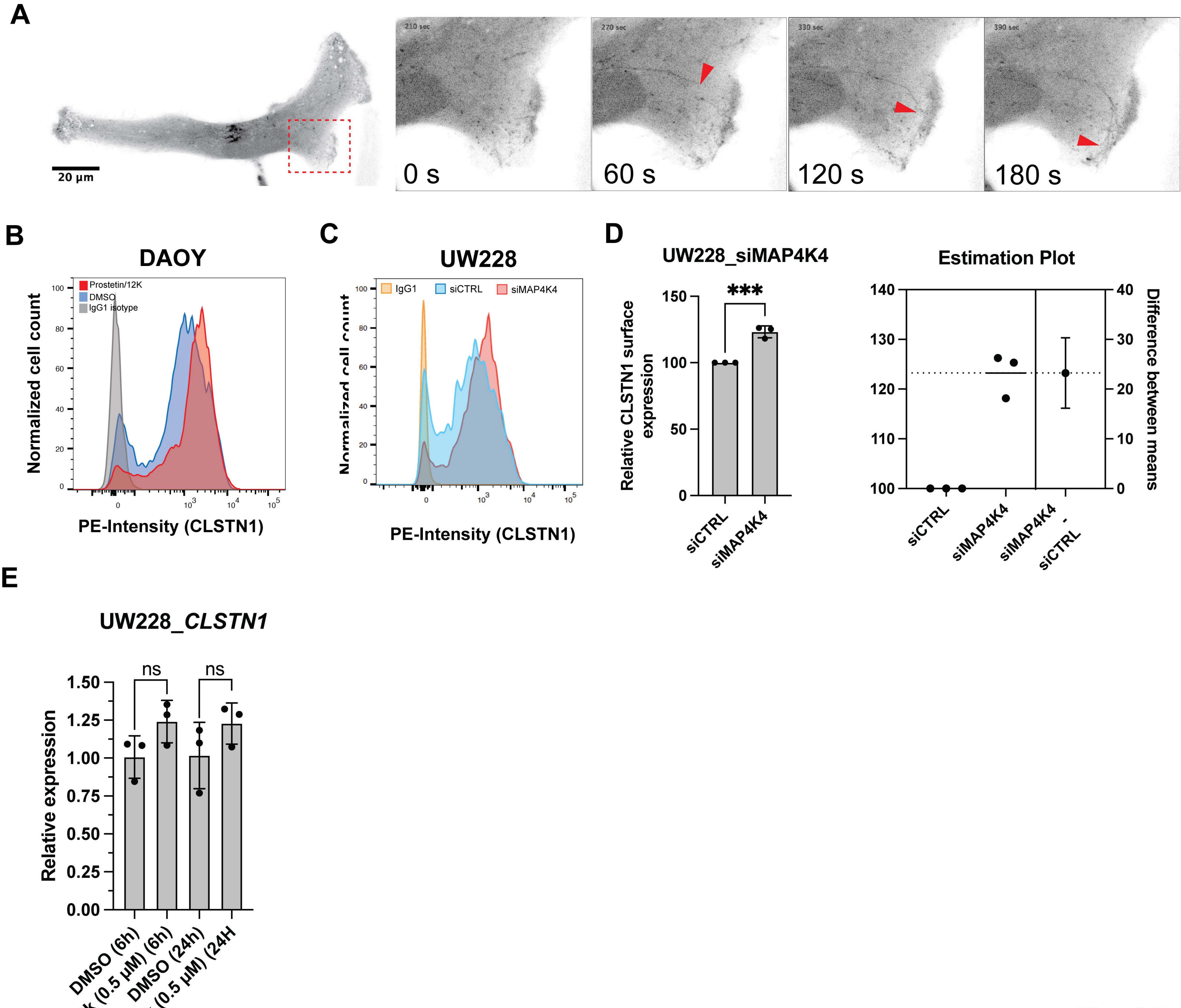

A
20 µm
210 sec
270 sec
330 sec
390 sec
0 s
60 s
120 s
180 s
B
DAOY
Prostetin/12K
DMSO
IgG1 isotype
Normalized cell count
PE-Intensity (CLSTN1)
C
UW228
IgG1
siCTRL
siMAP4K4
PE-Intensity (CLSTN1)
D
UW228_siMAP4K4
***
Relative CLSTN1 surface expression
siCTRL
siMAP4K4
Estimation Plot
Difference between means
siCTRL
siMAP4K4
siMAP4K4 - siCTRL
E
UW228_CLSTN1
ns
ns
Relative expression
DMSO (6h)
P/12k (0.5 µM) (6h)
DMSO (24h)
P/12k (0.5 µM) (24H

Fig. S5

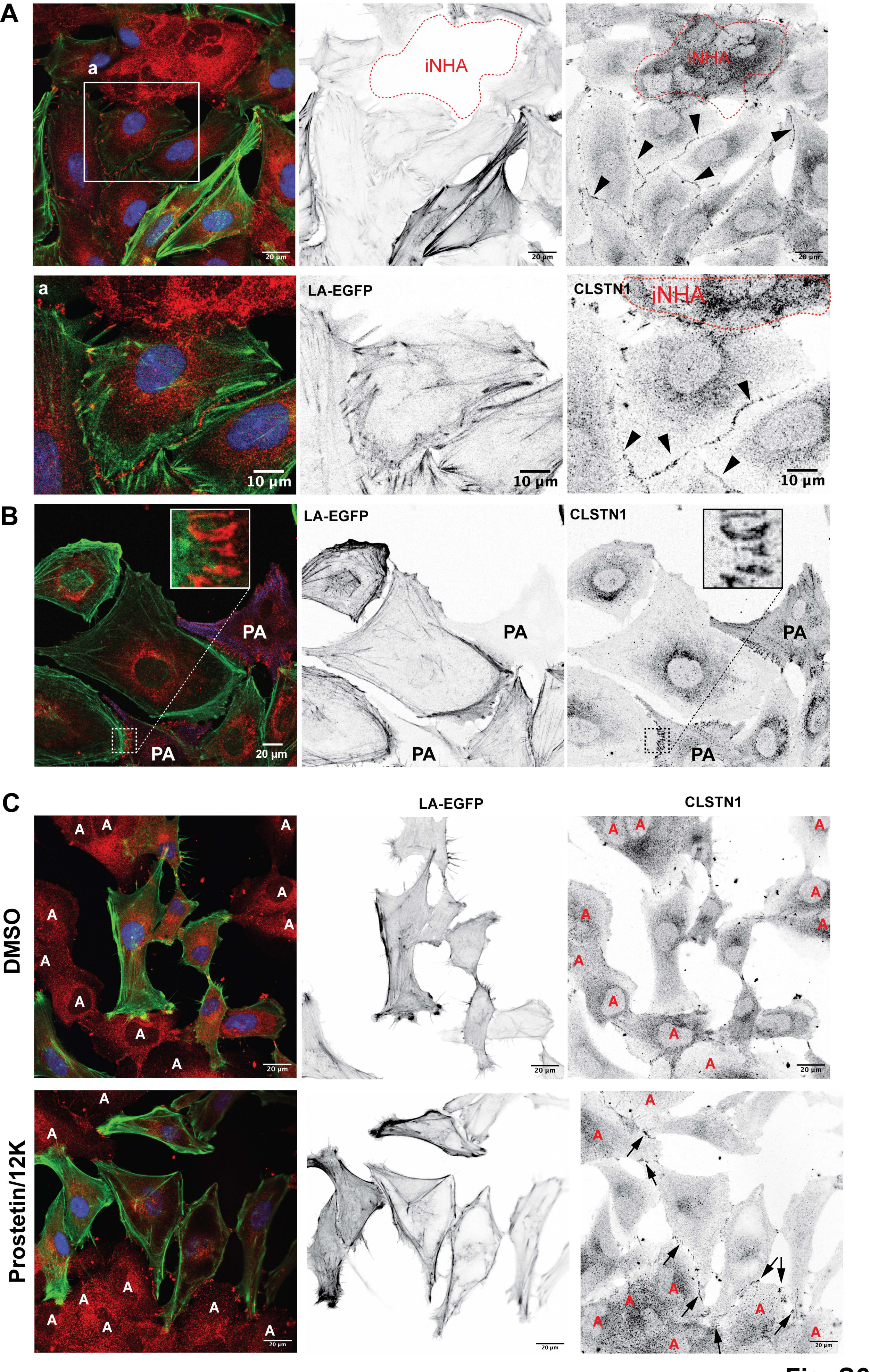

A
a
iNHA
20 µm
LA-EGFP
CLSTN1
10 µm
B
PA
20 µm
C
DMSO
Prostetin/12K

Fig. S6

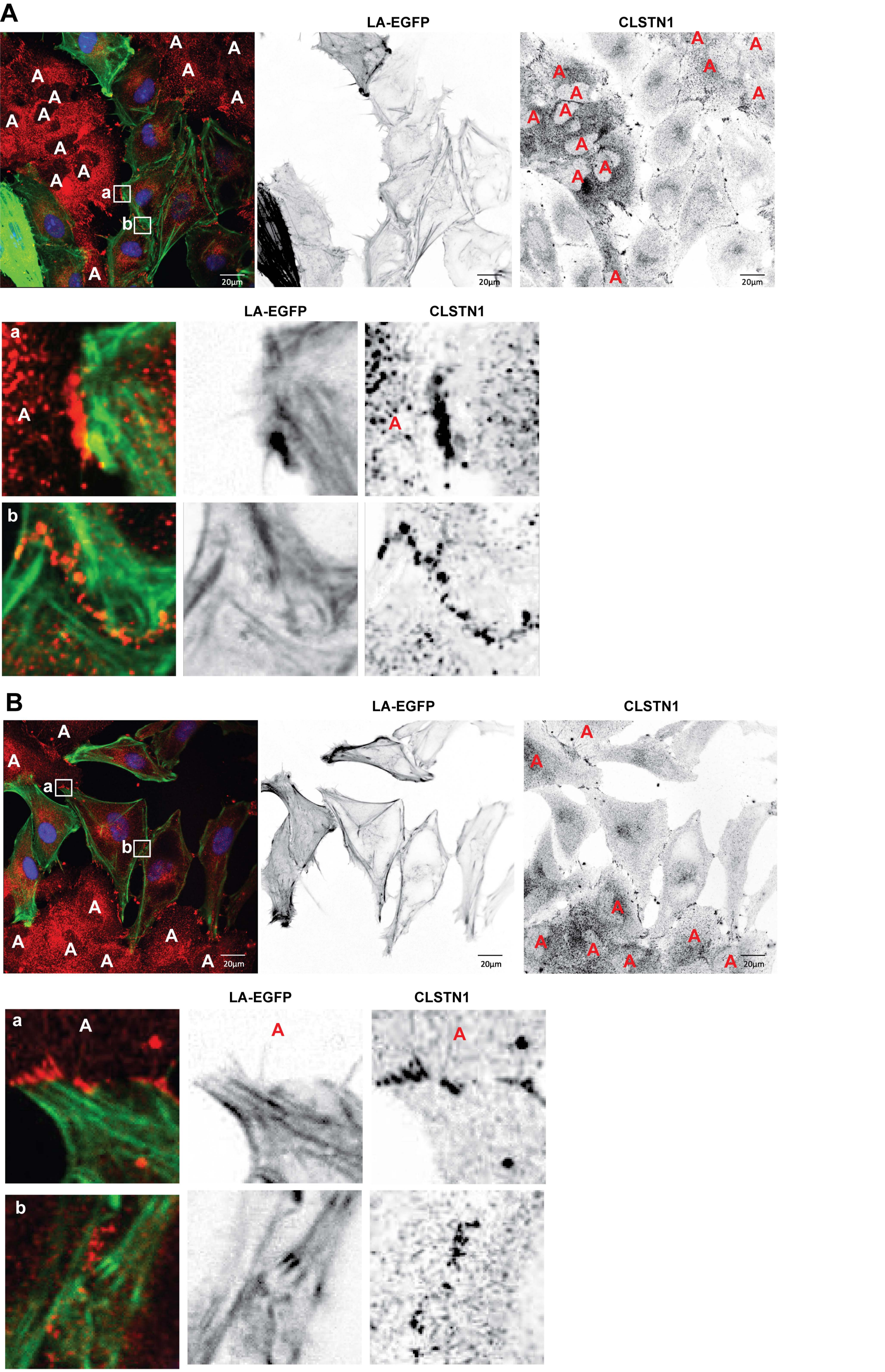

Fig. S7

Supplementary methods file

***Table 1:*** *Cell lines used*

| Name | Origin | Authentication/testing | Medium |
|---|---|---|---|
| ONS-76 | Michael Taylor lab, Sickkids Toronto, Canada | Single Nucleotide Polymorphism (SNP) typing, 02/2022)<br>Regular LookOut® Mycoplasma PCR Detection kit (#MP0035, Sigma Aldrich) | RMPI-1640 medium (R0883, Sigma) supplemented with GlutaMax (#35050-038, Gibco), 10% FBS (S0615, Sigma) and 1% penicillin/streptomycin (#15140-122, Gibco) |
| HD-MB03 | Till Milde, DKFZ Heidelberg, Germany | Single Nucleotide Polymorphism (SNP) typing, 02/2022)<br>Regular LookOut® Mycoplasma PCR Detection kit (#MP0035, Sigma Aldrich) | RMPI-1640 medium (R0883, Sigma) supplemented with GlutaMax (#35050-038, Gibco), 10% FBS (S0615, Sigma) and 1% penicillin/streptomycin (#15140-122, Gibco) |
| UW228 | John Silber, Seattle, USA | Single Nucleotide Polymorphism (SNP) typing, 02/2022)<br>Regular LookOut® Mycoplasma PCR Detection kit (#MP0035, Sigma Aldrich) | DMEM medium (Gibco, 11965092) supplemented with GlutaMax (#35050-038, Gibco), 10% FBS (S0615, Sigma) and 1% penicillin/streptomycin (#15140-122, Gibco) |
| D425-Med (D425) | Henry Friedman lab, Duke University, UK | Single Nucleotide Polymorphism (SNP) typing, 02/2022)<br>Regular LookOut® Mycoplasma PCR Detection kit (#MP0035, Sigma Aldrich) | IMEM medium (#10373-017, Gibco) supplemented with GlutaMax (#35050-038, Gibco), 10% FBS (S0615, Sigma) and 1% penicillin/streptomycin (#15140-122, Gibco) |
| D283-Med (D283) | ATTC | Single Nucleotide Polymorphism (SNP) typing, 02/2022)<br>Regular LookOut® Mycoplasma PCR Detection kit (#MP0035, Sigma Aldrich) | IMEM medium (#10373-017, Gibco) supplemented with GlutaMax (#35050-038, Gibco), 10% FBS (S0615, Sigma) and 1% penicillin/streptomycin (#15140-122, Gibco) |

***Table 2****: Primary antibodies used in Western Blot*

| Antibody | Source | Dilution | Company | Catalog Number |
|---|---|---|---|---|
| CLSTN1 | Rabbit | 1:1000 | Abcam | 134130 |
| N-cadherin | Rabbit | 1:1000 | Cell Signaling Technology | 13116T |
| Integrin-α-5 | Rabbit | 1:1000 | Cell Signaling Technology | 4705 |
| GAPDH | Rabbit | 1:1000 | Cell Signaling Technology | 2118L |
| Histone H3 | Rabbit | 1:1000 | Cell Signaling Technology | 44995 |
| Vimentin | Rabbit | 1:1000 | Cell Signaling Technology | 57415 |
| C-Jun | Rabbit | 1:1000 | Cell Signaling Technology | 9165 |

***Table 3:*** *Primary antibodies used in FACS*

| Antibody | Source | Dilution | Company | Catalog Number |
|---|---|---|---|---|
| CLSTN1 (Biotin-conjugated) | Rabbit | 1:100 | ABIN | 6876422 |
| IgG Isotype Control (PE-conjugated) | Rabbit | 1:300 | Cell Signaling Technology | 5742 |

***Table 4:*** *Secondary antibodies used in IFA*

| Antibody | Source | Dilution | Company | Catalog Number |
|---|---|---|---|---|

| | | | | |
|---|---|---|---|---|
| CLSTN1 | Rabbit | 1:100 | Abcam | 134130 |
| Connexin 43 (Cx43) | Rabbit | 1:1000 | Milipore | C6219 |
| Afadin 6 (AF-6) | Rabbit | 1:1000 | Millipore | C6219 |
| N-Cadherin | Rabbit | 1:1000 | Thermo Fisher Scientific | PA5-83130 |
| N-cadherin | Rabbit | 1:400 | Cell Signaling Technology | 13116T |
| GFAP | Goat | 1:250 | Abacam | Ab53554 |

***Table 5:*** *Secondary Antibodies Used in Western Blot*

| Antibody | Source | Dilution | Company | Catalog Number |
|---|---|---|---|---|
| Anti-rabbit IgG, HRP-linked | Goat | 1:5000 | Cell Signaling Technology | 7074S |
| Anti-mouse IgG, HRP-linked | Horse | 1:5000 | Cell Signaling Technology | 7076S |

***Table 6****: Secondary antibodies used in FACS*

| Antibody | Source | Dilution | Company | Catalog Number |
|---|---|---|---|---|
| PE Streptavidin | Rabbit | 1:200 | BioLegend | 405203 |

***Table 7:*** *Secondary antibodies used in IFA*

| Antibody | Source | Dilution | Company | Catalog Number |
|---|---|---|---|---|
| Alexa Fluor 647 – Anti-Rabbit IgG (H+L) | Donkey | 1:250 | Jackson ImmunoResearch | 711-605-152 |
| Alexa Fluor 421- Anti-Goat IgG (H+L) | Donkey | 1:200 | Jackson ImmunoResearch | 705-675-147 |

Primary and secondary antibodies used in Western Blot, FACS, and IFA were diluted with 1X TBS-T 5% non-fat dry milk, 2% FBS in PBS, and 5% FBS in PBS, respectively.

***Table 7:*** *siRNAs*

| Target Gene | Company | Catalog Number |
|---|---|---|
| siCtrl (scrambled) | Dharmacon | D-001210-02-05 |
| CLSTN1 (si1-CLSTN1) | Dharmacon | J-020393-05-0002 |
| CLSTN1 (si2-CLSTN1) | Dharmacon | J-020393-06-0002 |
| CLSTN1 (si3-CLSTN1) | Dharmacon | J-020393-07-0002 |
| CLSTN1 (si4-CLSTN1) | Dharmacon | J-020393-08-0002 |
| MAP4K4 (siMAP4K4) | | |

***Table 8:*** *Lentiviral Vectors*

| Vector Name | Vector ID | Company | Selection Marker |
|---|---|---|---|
| pLV[Exp]-Bsd-CMV>hCLSTN1[NM_001009566.3](ns):3xGGGGS: | *VB230510-1232gzf* | VectorBuilder | Bsd (blasticidin resistance gene) |

| | | | |
|---|---|---|---|
| mNeonGreen | | | |
| pLV[Exp]-Bsd-CMV>hCLSTN1[NM_001009566.3]/3xGGGGS | *VB230510-1225fgu* /V5. | VectorBuilder | Bsd (blasticidin resistance gene) |

CLSTN1-coding sequences in lentiviral vectors:

**pLV_Bsd-CMV_hCLSTN1_3xGGGGS_V5:**

```
MGGRRVRWEVYISRAGLVNRQIQVCTKKQAATMLRRPAPALAPAARLLLAGLLCGGGVWAARVNKHKPWLEPTYH
GIVTENDNTVLLDPPLIALDKDAPLRFAESFEVTVTKEGEICGFKIHGQNVPFDAVVVDKSTGEGVIRSKEKLDC
ELQKDYSFTIQAYDCGKGPDGTNVKKSHKATVHIQVNDVNEYAPVFKEKSYKATVIEGKQYDSILRVEAVDADCS
PQFSQICSYEIITPDVPFTVDKDGYIKNTEKLNYGKEHQYKLTVTAYDCGKKRATEDVLVKISIKPTCTPGWQGW
NNRIEYEPGTGALAVFPNIHLETCDEPVASVQATVELETSHIGKGCDRDTYSEKSLHRLCGAAAGTAELLPSPSG
SLNWTMGLPTDNGHDSDQVFEFNGTQAVRIPDGVVSVSPKEPFTISVWMRHGPFGRKKETILCSSDKTDMNRHHY
SLYVHGCRLIFLFRQDPSEEKKYRPAEFHWKLNQVCDEEWHHYVLNVEFPSVTLYVDGTSHEPFSVTEDYPLHPS
KIETQLVVGACWQEFSGVENDNETEPVTVASAGGDLHMTQFFRGNLAGLTLRSGKLADKKVIDCLYTCKEGLDLQ
VLEDSGRGVQIQAHPSQLVLTLEGEDLGELDKAMQHISYLNSRQFPTPGIRRLKITSTIKCFNEATCISVPPVDG
YVMVLQPEEPKISLSGVHHFARAASEFESSEGVFLFPELRIISTITREVEPEGDGAEDPTVQESLVSEEIVHDLD
TCEVTVEGEELNHEQESLEVDMARLQQKGIEVSSSELGMTFTGVDTMASYEEVLHLLRYRNWHARSLLDRKFKLI
CSELNGRYISNEFKVEVNVIHTANPMEHANHMAAQPQFVHPEHRSFVDLSGHNLANPHPFAVVPSTATVVIVVCV
SFLVFMIILGVFRIRAAHRRTMRDQDTGKENEMDWDDSALTITVNPMETYEDQHSSEEEEEEEEEEEESEDGEEED
DITSAESESSEEEEGEQGDPQNATRQQQLEWDDSTLSYSGGGGSGGGGSGGGGSGKPIPNPLLGLDST
```

**VB230510_pLV_Bsd_hCLSTN1_3xGGGGS_mNeonGreen**

```
MGGRRVRWEVYISRAGLVNRQIQVCTKKQAATMLRRPAPALAPAARLLLAGLLCGGGVWAARVNKHKPWLEPTYH
GIVTENDNTVLLDPPLIALDKDAPLRFAESFEVTVTKEGEICGFKIHGQNVPFDAVVVDKSTGEGVIRSKEKLDC
ELQKDYSFTIQAYDCGKGPDGTNVKKSHKATVHIQVNDVNEYAPVFKEKSYKATVIEGKQYDSILRVEAVDADCS
PQFSQICSYEIITPDVPFTVDKDGYIKNTEKLNYGKEHQYKLTVTAYDCGKKRATEDVLVKISIKPTCTPGWQGW
NNRIEYEPGTGALAVFPNIHLETCDEPVASVQATVELETSHIGKGCDRDTYSEKSLHRLCGAAAGTAELLPSPSG
SLNWTMGLPTDNGHDSDQVFEFNGTQAVRIPDGVVSVSPKEPFTISVWMRHGPFGRKKETILCSSDKTDMNRHHY
SLYVHGCRLIFLFRQDPSEEKKYRPAEFHWKLNQVCDEEWHHYVLNVEFPSVTLYVDGTSHEPFSVTEDYPLHPS
KIETQLVVGACWQEFSGVENDNETEPVTVASAGGDLHMTQFFRGNLAGLTLRSGKLADKKVIDCLYTCKEGLDLQ
VLEDSGRGVQIQAHPSQLVLTLEGEDLGELDKAMQHISYLNSRQFPTPGIRRLKITSTIKCFNEATCISVPPVDG
YVMVLQPEEPKISLSGVHHFARAASEFESSEGVFLFPELRIISTITREVEPEGDGAEDPTVQESLVSEEIVHDLD
TCEVTVEGEELNHEQESLEVDMARLQQKGIEVSSSELGMTFTGVDTMASYEEVLHLLRYRNWHARSLLDRKFKLI
CSELNGRYISNEFKVEVNVIHTANPMEHANHMAAQPQFVHPEHRSFVDLSGHNLANPHPFAVVPSTATVVIVVCV
SFLVFMIILGVFRIRAAHRRTMRDQDTGKENEMDWDDSALTITVNPMETYEDQHSSEEEEEEEEEEEESEDGEEED
DITSAESESSEEEEGEQGDPQNATRQQQLEWDDSTLSYSGGGGSGGGGSGGGGSMVSKGEEDNMASLPATHELHI
FGSINGVDFDMVGQGTGNPNDGYEELNLKSTKGDLQFSPWILVPHIGYGFHQYLPYPDGMSPFQAAMVDGSGYQV
HRTMQFEDGASLTVNYRYTYEGSHIKGEAQVKGTGFPADGPVMTNSLTAADWCRSKKTYPNDKTIIS
```
